# Developmental co-emergence of cardiac and gut tissues modeled by human iPSC-derived organoids

**DOI:** 10.1101/2020.04.30.071472

**Authors:** A.C. Silva, O.B. Matthys, D.A. Joy, M.A. Kauss, V. Natarajan, M.H. Lai, D. Turaga, A.P. Blair, M. Alexanian, B.G. Bruneau, T.C. McDevitt

## Abstract

During embryogenesis, paracrine signaling between tissues in close proximity contributes to the determination of their respective cell fate(s) and development into functional organs. Organoids are *in vitro* models that mimic organ formation and cellular heterogeneity, but lack the paracrine input of surrounding tissues. Here, we describe a human multilineage iPSC-derived organoid that recapitulates cooperative cardiac and gut development and displays extensive cellular and structural complexity of both tissues. We demonstrate that the presence of endoderm tissue (gut/intestine) in multilineage organoids contributed to the development of the cardiac tissue, specifically cardiomyocyte expansion, compartmentalization, enrichment of atrial/nodal cells, myocardial compaction and functional fetal-like maturation. Overall, this study demonstrates the ability to generate specific cooperative tissues originating from different germ lineages within a single organoid model, an advance that will further the examination of multi-tissue interactions during development and disease.

## Introduction

The formation of organs during embryogenesis results from a coordinated sequence of morphogenic events that regulate the co-emergence and self-organization of distinct cell populations (Zhang and Hiiragi, 2018). Organoids derived from stem and progenitor cells provide an experimentally tractable model of tissue morphogenesis *in vitro* due to the fact that they differentiate into multiple cell types in a spatially-organized manner and recapitulate functional properties of developing native tissues (Lancaster and Knoblich, 2014;Matthys *et al.*, 2020). However, most organoids to date give rise to single tissues in isolation, and lack the mechanical cues and paracrine signaling contributed by neighboring organs that synergistically benefit the trajectory of tissue development *in vivo* (Varner and Taber, 2012;Jung *et al.*, 1999;Ishii *et al.*, 2007). Previous attempts to generate multi-tissue organoids in a controllable fashion have mainly involved the fusion of pre-differentiated single-lineage organoids (Koike *et al.*, 2019;Bagley *et al.*, 2017).

Mesoderm and endoderm display reciprocal interactions during embryonic development that are fundamental to the successful formation of several organs, including the heart (Lough and Sugi, 2000;Schultheiss *et al.*, 1995;Sugi and Lough, 1994). *In vitro* co-culture systems of embryonic germ layer explants and pluripotent stem cells demonstrate that endoderm cells secret cardiac-inductive factors (e.g. Activin A, FGF2) that promote cardiomyocyte formation (Anderson *et al.*, 2016;Rudy-Reil and Lough, 2004;Brown *et al.*, 2010;Uosaki *et al.*, 2012;Sugi and Lough, 1995). However, it has been challenging to determine the effect and sequence of direct interactions between endoderm and mesoderm *in vivo* because interrogation at each stage of development disrupts the natural course of morphogenesis.

Here, we generated 3D spheroids of early committed mesendoderm progenitors in a culture medium permissive of multi-lineage differentiation. We observed the co-emergence of mesoderm and endoderm cell lineages, resulting in unique multilineage organoids able to spontaneously organize the formation of complex structures similar to the developing heart and gut. Multiple *in vitro* models have been developed to study various aspects of the heart, including patterning constructs (Ma *et al.*, 2015), gastruloids/organoids (Lee *et al.*, 2020;Rossi *et al.*, 2020),(Drakhlis *et al.*, 2021) (development), and engineered microtissues (homeostasis and maturation) (Ronaldson-Bouchard *et al.*, 2018;Hookway *et al.*, 2019;Mills *et al.*, 2017;Macqueen *et al.*, 2018). However, models that recapitulate heart morphogenic dynamics (i.e. gastruloid/ organoids) have been mostly obtained with mouse stem cells (Rossi *et al.*, 2020;Lee *et al.*, 2020), and only recently using human induced pluripotent stem cells (iPSCs)(Drakhlis *et al.*, 2021). On the other hand, intestinal tissue has been generated *in vitro* with high reproducibility using mouse and human stem cells derived from tissue biopsies and iPSC (Sato *et al.*, 2009;Munera *et al.*, 2017). The cardiac and gut (multilineage) organoid presented here constitutes a first step toward a new generation of organoids that recapitulate the co-development and differentiation of two defined tissues for long periods of time (>1 year). Such defined multi-lineage/multi-organ organoids hold the unique promise of modeling the crosstalk between specific tissues that occurs during human embryogenesis, tissue maturation and disease— a complex phenomenon that is difficult to dissect *in vivo*.

## Results

### Formation of heart-like compartments in multilineage spheroids

We began by generating spheroids from mesendoderm progenitors differentiated for 5 days with the GiWi protocol (Lian *et al.*, 2012). Spheroids were then cultured in conventional, cardiac-permissive medium supplemented with insulin (RPMI/B27+), or in an ascorbic acid (AA)-supplemented medium formulated for epicardial cell differentiation to mimic the signaling that occurs during cardiac differentiation to support the emergence of a variety of cell types (i.e. fibroblasts and smooth muscle cells (SMC))(Bao *et al.*, 2016;Lian *et al.*, 2013) **(Figure 1A,B)**. Noticeable differences in size and structural arrangement between spheroids grown in conventional versus AA medium (hereafter referred to as “conventional” and “multilineage” microtissues, respectively) became evident within 2 weeks of differentiation, with multilineage microtissues growing to approximately 3 times the size of conventional microtissues by 30 days **(Figure 1C, D, E)**. After 10 days, conventional microtissues displayed a spherical, uniformly translucent morphology with spontaneous contractility throughout **(Figure 1C, D; Movie S1)**, whereas the multilineage microtissues exhibited a dark core of vigorously contracting cells surrounded by translucent non-contracting cells. Calcium transients were limited to the central region of contracting cells in the multilineage tissues **(Figure 1C,D; Movie S2)**. At early stages of culture, light-sheet fluorescence microscopy revealed that conventional microtissues were largely composed of cardiomyocytes, as identified by cardiac troponin (cTnT) immunostaining, that co-expressed TBX18 (cTnT+/TBX18+ cells), and an outer layer of TBX18+ epicardial cells (cTnT-/TBX18+) **(Figure 1F; S1A, S2A)**. However, by day 40, conventional microtissues had only small patches of epicardial cells at the surface **(Figure 1F; S2A)**. In contrast, multilineage microtissues displayed a core of cTnT+ cardiomyocytes surrounded by a non-myocyte, stromal-like cell population (phalloidin+, cTnT-), and an outer layer of TBX18+ epicardial cells (cTnT-/TBX18+), which was preserved for more than two month **(Figure 1F; S1A, S2A)**, and better resembled the radial organization of the embryonic heart wall **(Figure S1A)**. Additional histological analysis revealed that both conventional and multilineage microtissues contained SMCs (SMA+/MYH7−; **Figure S2B**). However, in multilineage microtissues, cardiomyocytes positive for the ventricular marker MLC2v aggregated together in the cardiac tissue core, indicating spatial compartmentalization reminiscent of the distinct emergence of atrial and ventricular cardiomyocytes in the primitive heart **(Figure 1G)**; distinct spatial compartmentalization of cardiomyocytes was not observed in conventional microtissues.

**Figure 1.**
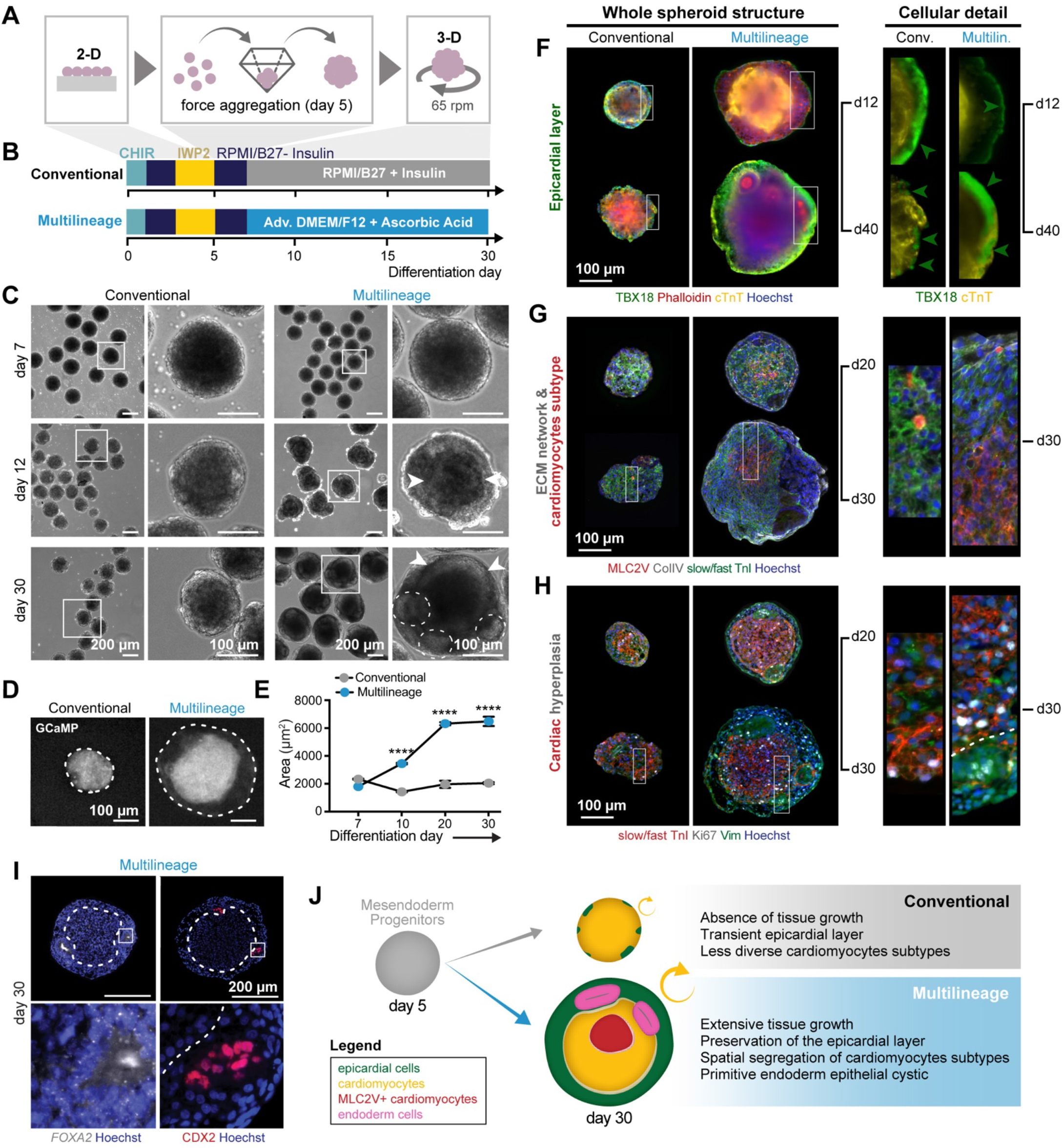
Derivation of a multilineage organoid with complex cardiac structures. **A-B** Overall schematic of conventional and multilineage microtissue formation. **C.** Brightfield images of conventional and multilineage microtissues over the course of 30 days in cell culture; high magnification views of the regions within white squares (right). Multilineage microtissues develop a compact dark core of cells (arrowheads) and epithelial-like structures (dashed outlines) at the surrounding translucent region. **D.** Calcium flux transients visualized by GCAMP are limited to the center area of the multilineage microtissues, whereas they occur throughout the conventional microtissues. Dashed line: microtissue boundaries. **E.** Multilineage microtissues increase in surface area throughout a 30-day culture. Representative experiment. Conventional vs. Multilineage: ****p<0.0001. **F-I.** Structural dynamics and cell composition of conventional and multilineage organoids over the course of 30-40 days in culture. **F.** Representative light-sheet imaging of microtissues stained for cardiac Troponin T (cTnT) and TBX18. Conventional microtissues are composed of cardiomyocytes positive for cTnT covered by a continuous outer layer of epicardial-like cells (cTnT-/TBX18+). However, the epicardial coverage shrinks over time to small patches of cells. Multilineage microtissues contain a core of cardiomyocytes (cTnT+/TBX18-) surrounded by stromal-like cells positive for phalloidin and a continuous outer layer of epicardial-like cells (cTnT-/TBX18+). Like in conventional microtissues, the epicardial layer of the multilineage microtissues suffers some disruption throughout the course of the culture, but overall is better preserved over time. Arrowhead: epicardial layer. **G.** Multilineage microtissues evidence a collagen IV network and MYL2+ cardiomyocytes that cluster at the center of the microtissues. Conventional microtissues demonstrate a network of collagen IV at day 20 (d20), but it was not evident at later stages. No MYL2+ positives cells were detected in conventional microtissues. **H.** Multilineage microtissues exhibit more actively cycling (ki67+) cardiomyocytes and non-cardiomyocytes than conventional microtissues. **I.** At day 30, multilineage organoids develop epithelial-like structures positive for mid-hindgut markers (*FOXA2*/CDX2). **J.** Schematic of the main contrasting features between conventional and multilineage microtissues. Due the fact multilineage microtissues contained epithelial endoderm, displayed greater cardiac structural complexity, and exhibited extensive growth dynamics, they were hereafter referred to as multilineage organoids.

The dramatic growth of multilineage microtissues over the first 30 days in culture was paralleled by an increase in the volume of the cardiac region **(Figure 1H; S1C, S2C; Movie S3-4)**. Cardiac growth appeared to result from cardiomyocyte proliferation, based upon immunostaining for Ki67, a marker of active cell cycle—analogous to cardiac wall thickening during development **(Figure 1H; S1C)**. In addition, at day 30, we observed the emergence of small epithelial cystic structures within the previously translucent layers of cells enclosing the cardiac core in multilineage microtissues **(Figure 1C)**. These cystic structures expressed the mid-hindgut (MHG) endoderm markers FOXA2 and CDX2 **(Figure 1C,I; S2D,E)**. Since the multilineage microtissues contained endoderm epithelial clusters, attained greater structural complexity typical of the cardiac compartment and exhibited growth dynamics comparable to the embryonic heart, they were hereafter referred to as multilineage organoids **(Figure 1J)** (Lancaster and Knoblich, 2014).

### Long-term culture of multilineage organoids recapitulates embryonic heart and gut development

We next examined whether multilineage organoids acquired further complexity with extended time in culture. The growth of multilineage organoids accelerated throughout culture duration, with the average organoid reaching ~2mm in diameter after 100 days **(Figure 2A-C)**. After 50 days, the stromal cells surrounding the cardiac core were replaced by a mesenchymal tissue **(Figure 2D)** enriched in extracellular matrix (ECM) components, such as hyaluronic acid **(Figure S2F),** and a fine fibrillar network of glycosaminoglycans **(Figure 2D, light green/blue)** that was populated by a few *WT1*-expressing cells **(Figure S1A)**. The mesenchymal tissue resembled the interstitial tissue underlying the epicardium (or sub-epicardium) and the peritoneum (protective layer of the intestine) **(Figure 2D)**. In addition, smooth muscle actin (SMA)-positive cells were detected at the surface of the organoid and in the sub-epicardium, and a few endothelial cells (PECAM-1+) appeared interspersed among the cardiomyocytes **(Figure 2F)**. By day 60, SMCs (αSMA+ MYH7-) emerged prominently between the cardiac and the primitive gut-like tissue. The SMC tissue displayed slow and infrequent calcium transients (~5s duration per transient; 1-3 transients/minute) that corresponded to large contractions (>100 μm) that resembled peristalsis **(Figure 2G,H; Movie S5-6)**. More endothelial cells were detected between the cardiomyocytes, forming branched-like structures resembling pre-vascular networks **(Figure 2F)**. Between 70 and 100 days of differentiation, the primitive gut tissues matured, as shown by the development of Paneth cells (antimicrobial peptide-producing cells) and goblet cells (which produce mucins) **(Figure 2D)**. Thus, multilineage organoids undergo a sequence of morphogenic events that recapitulates the development of the human embryonic heart and gut, including the initial emergence of cardiac tissue followed by gut tissue, with interstitial tissue (sub-epicardium/sub-peritoneum-like tissue) separating the two distinct tissues.

**Figure 2.**
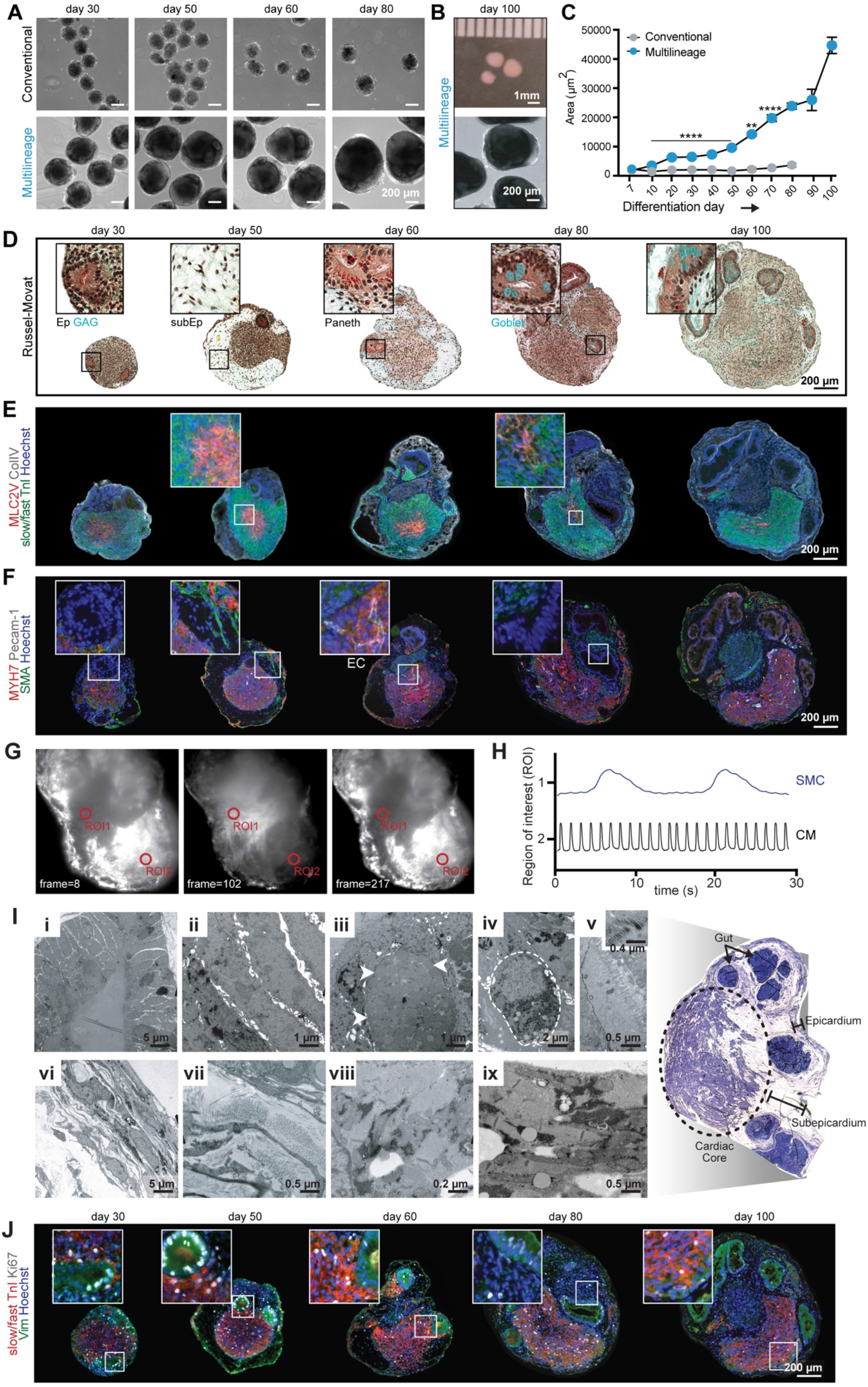
Long-term culture of multilineage organoids recapitulates early heart and gut development. **A.** Brightfield microscopy images of conventional and multilineage tissues over days 30 to 80 of culture. Conventional microtissues are viable until day 80 and beyond, whereas conventional microtissues degenerate. **B.** Stereoscope macroscopy (top) and brightfield (bottom) images of day-100 multilineage organoids. **C.** Multilineage organoids display an increase in surface area until day 100 of culture. Representative experiment. Conventional vs. Multilineage: **p<0.005, ****p<0.0001. **D-F.** Structural dynamics and cell composition of multilineage organoids over the course of the culture as revealed by Russel-Movat stain (D) and fluorescent immunostaining (E-F). **G-H.** Representative images (G) and traces (H) of spontaneous calcium flux transients of the smooth muscle cells (ROI 1) and cardiomyocytes (ROI 2) of a multilineage organoid imaged via light sheet microscopy (Movie S6). **I.** Transmission electron microscopy of the gut structures (i-v), epicardium/subepicardium (vi-vii) and cardiac core (viii-ix). GAG, glycosaminoglycans; Ep, epithelial-like structures; subEp, subepicardium-like tissue; Endo, endothelial cells network; Arrowheads, goblet cell (iii); Dashed line, Paneth cell (iv). **J.** The number of actively cycling cells (Ki67+) decreases over time in multilineage organoids.

To assess the cellular heterogeneity of the multilineage organoids, we performed single-cell RNA-sequencing (scRNA-seq) on dissociated organoids after 100 days. Seurat analysis revealed 6 cell clusters corresponding to cardiomyocytes, SMCs, proliferating cells, fibroblasts, gut and enteroendocrine cells **(Figure 3A-C; S3A; Table S1)**. GO term analysis of the enteroendocrine cell cluster yielded hormone transport and secretion ( expressing CHGA; **Figure 3B; S3A**), suggesting the development of chemosensing properties in the gut compartment(Gribble and Reimann, 2016). Sub-clustering of the gut cells confirmed the differentiation of specialized intestinal cells, including: 1) enterocyte-like cells (absorptive cells expressing VIL1, ANPEP, MUC13/17); 2) Paneth cells (expressing LYZ); 3) LGR5-expressing intestinal stem cells; and 4) small intestine cells marked by high expression of APOB, CDX2 and VIL1, along with enrichment of genes for glucose and cholesterol transporters (SLC2A2 and NPC1L1, respectively), and digestion (FABP2, SI, SLC5A1) **(Figure 3C,D; S3A,B)**. The presence of these gut cell types was further validated by electron microscopy of 100-day old multilineage organoids through the identification of key structural features of each cell type **(Figure 2I i-v)**. Goblet cells were not detected by scRNA-seq, presumably due to their larger size.

**Figure 3.**
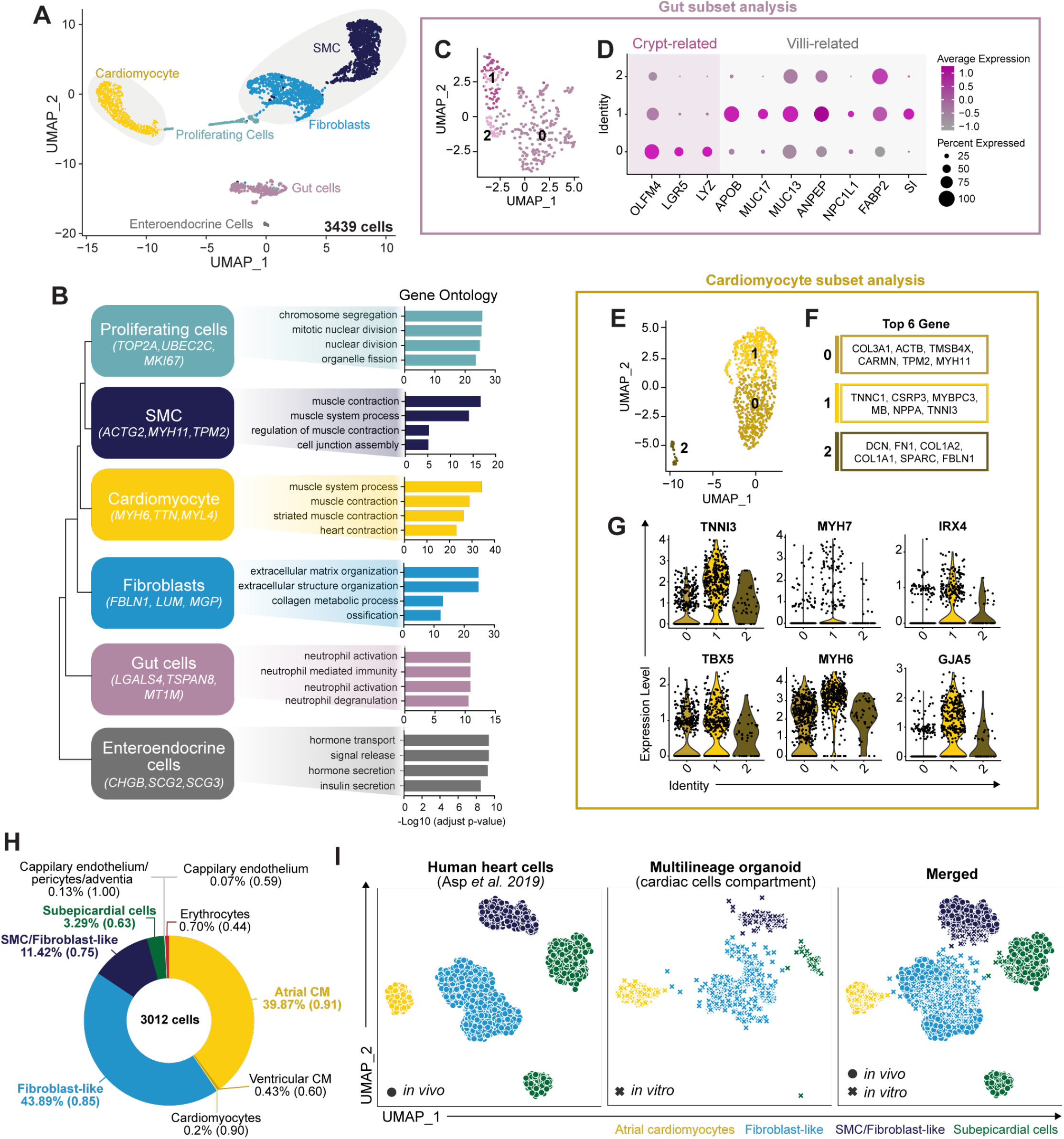
Single-cell transcriptomic analysis at day 100 reveals complex cellular heterogeneity of multilineage organoids. **A.** UMAP plot of all cellular populations identified. Light gray: Seurat cell clusters analyzed using an *in vivo* heart cells classifier. **B.** Cluster tree analysis of the cell clusters with top differentially expressed genes defining each cluster (left) and gene ontology analysis of associated biological processes (right). **C-D.** A sub-clustering analysis of the gut cells evidenced 3 main cell clusters (C) distinguished by the expression of crypt-related or villi-related genes (D): cluster 0 composed mainly of crypt cells (OLFM4), such as gut stem cells (LGR5) and Paneth cells (LYZ); and cluster 1 and 2, constituted by cells at the villi, the enterocytes, identified by the expression of genes related to transmembrane mucins (MUC13, MUC17), digestion (ANPEP, FABP2, SI) and cholesterol transporter genes (NPC1L1), as well as APOB, a gene expressed in the small intestine. **E-F.** Subset analysis of the cardiomyocyte cluster (E) and top-6 genes of the three sub-clusters (F) identify clusters 0 and 2, composed of more immature cardiomyocytes, and cluster 1, composed of more mature cells. **G.** Violin plots of secondary cardiomyocyte analysis depicting expression of key cardiomyocyte subtype genes for ventricular-(top) and atrial-like cells (bottom). **H.** Identity of the cardiac-like cells of multilineage organoids, as determined by a cell classifier trained on expression data from human fetal heart cells. The classifier was applied to the multilineage organoid cardiomyocyte, fibroblast and smooth muscle cells (SMC) clusters identified by Seurat analysis. Data displayed include cell percentage and assignment probability of transcriptomic resemblance with *in vivo* heart cells. **I.** UMAP plot of *in vivo* human developing heart cells and *in vitro* multilineage organoid cells classified as atrial cardiomyocyte, subepicardial cells, fibroblast-like, smooth muscle cells/fibroblast-like cells (probability greater or equal to 0.85). The UMAP projection demonstrates that *in vitro* multilineage organoid cells align closely to the *in vivo* human heart cells suggesting that the transcriptome of the multilineage organoid heart cells closely mimic that of human developing heart cells, and favors atrial cardiomyocyte specification.

Analysis of cell populations within the cluster enriched for proliferative markers revealed the presence of proliferating cardiomyocytes (TTN, MYH6, NPPA) and SMCs (ACTG2, MYH11) **(Figure S3C,D)**. Ki67 immunostaining demonstrated a decline in the number of actively cycling cells over time **(Figure 2J)**; nevertheless, some cardiomyocytes (slow/fast TnI+), SMCs, and gut cells remained Ki67+ after 100 days in culture, corroborating the scRNA-seq results **(Figure S3G)**. Sub-clustering analysis of the cardiomyocytes indicated that after 100 days in culture, cardiomyocytes were divided into 3 groups, separated mainly by differences in maturation stage markers: immature cells that either expressed ECM related genes(Cui *et al.*, 2019) (Cluster 2) or smooth muscle myosin (MYH11, Cluster 0), and more mature-like cardiomyocytes (Cluster 1) that expressed higher levels of TNNI3 **(Figure 3E,F)**. A minority of cardiomyocytes (~30%) expressed the ventricular cardiomyocyte markers IRX4 or MYH7 **(Figure 3G; 3SA)**, indicating that multilineage organoids favor the differentiation and maturation of atrial cardiomyocytes, as confirmed by the high expression of atrial markers such as TBX5, MYH6 and GJA5 **(Figure 3G)**. Although key atrial and ventricular genes were expressed in the same cluster of cells (Cluster 1), these genes were not co-expressed in individual cells **(Figure S3F)**.

To identify *in vivo* correlates, we used a cell type classifier, trained on scRNA-seq of human fetal heart tissue collected at 6.5-7 weeks **(Figure S3H)** (Asp *et al.*, 2019), to characterize a subset of cardiomyocyte, fibroblast and SMC clusters that were identified by default Seurat clustering and cardiac cell compartmentalization genes. The classifier estimated that 98% of the organoid cardiomyocytes were atrial cardiomyocytes (prediction probability = 0.91) **(Figure 3H, I; S3G)**, with classification driven by MYH6 expression, a classifier top feature for atrial cell predictions **(Table S2)**. Ventricular cells only represented 1% of the cardiomyocytes (prediction probability = 0.69) **(Figure 3H, I; S3G)**. The cell classifier also identified fibroblast and SMC/Fibroblast-like cells, as expected, which represented 44% and 11% of the cells analyzed with prediction probabilities of 0.85 and 0.75, respectively **(Figure 3H,I)**. In addition, the *in vivo* cell classifier assigned cell type identities not detected using default Seurat clustering analysis, such as sub-epicardial and capillary endothelium/pericytes/adventia cells, which together constituted less than 4% of the total cells evaluated **(Figure 3H,I; S3G)**. Although the capillary endothelium/pericytes/adventia cells corresponded to only 0.1% of the cells, their assignment had a high prediction probability of 1.00 **(Figure 3H; S3G)**. In summary, the *in vivo* cell classifier corroborated the cellular complexity of the multilineage organoids first observed by histological analysis, and suggested a high transcriptomic similarity between cells from multilineage organoids and cells from fetal human hearts **(Figure 3I)**.

Multilineage organoids remained viable for over 1 year in culture, and stably maintained their millimeter size-scale and structural complexity **(Figure S4)**. Contractile behavior was observed only when electrical stimulation was applied, a feature consistent with functionally mature iPSC-derived cardiomyocytes (Yang *et al.*, 2014) **(Figure S4B)**. The cardiac and intestine tissues remained localized at opposite ends of the organoids, separated by stromal cells and SMCs **(Figure S4C-E,G)**. In contrast to previous timepoints, small clusters of SMCs were detected amongst the cardiomyocytes in these aged organoids, and SMCs were more pronounced at the boundary of cardiac tissue regions **(Figure S4F)**. In addition, proliferating non-myocytes (Ki67+, cTnT-) were detected within the cardiac tissue, along with structures that resembled mature blood vessels (expressing lectin, a glycoprotein of the basal lamina of endothelial cells) embedded within or adjacent to the cardiac regions in some organoids **(Figure S4C, G)**. Organoids cultured for 1 year proved challenging to dissociate to single cells, potentially due to the structural complexity and ECM composition, thus limiting our ability to capture the morphological features and the transcriptional signature of individual cells. Overall, these observations demonstrate that multilineage organoids represent a new type of compartmentalized organoid model that exhibits the co-development and maturation of tissues derived from two distinct germ layers, heart (mesoderm) and gut (endoderm)(Koike *et al.*, 2019;Faustino Martins *et al.*, 2020;Bagley *et al.*, 2017).

### Multilineage organoids promote atrial/nodal cardiomyocyte specification and maturation

To further assess the maturity and sub-type specification of the cardiomyocytes, we performed electrophysiological and structural analysis of the intact spheroids and dissociated cells. Over time, 100% of conventional microtissues exhibited spontaneous contractility, and 33±21% of day-80 conventional microtissues could be paced at 2Hz **(Figure 4A; S5A,B)**. Furthermore, conventional microtissues exhibited no dramatic changes over time in kinetic calcium handling properties (amplitude and stroke velocities) under 1-Hz of stimulation, reflecting relatively immature function **(Figure 4B,C)**. In contrast, multilineage organoids demonstrated a decrease in spontaneous calcium activity over time, from 100% at day 10 to 37±16% at day 100, and an ability to respond to higher frequencies of electrical stimulation, up to 8Hz after 80 days of culture **(Figure 4A-D; S5A)**. In addition, after 30 days, multilineage organoids demonstrated progressively increasing amplitude and maximum stroke velocities under 1-Hz of stimulation, consistent with greater functional maturation than the conventional microtissues **(Figure 4B,C).**

**Figure 4.**
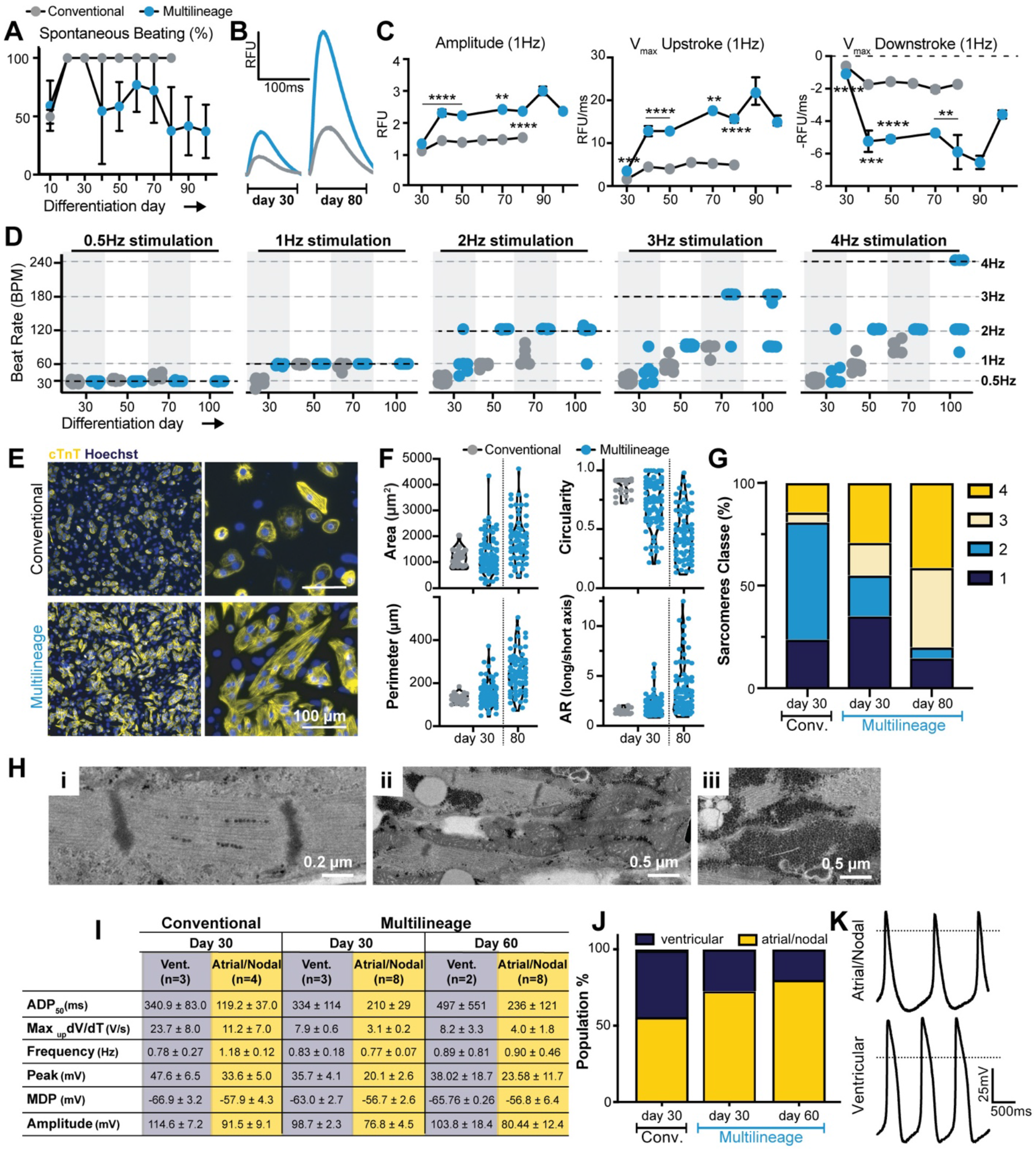
Cardiomyocytes from multilineage organoids display phenotypic and functional maturation throughout culture, and preferentially specify into atrial/nodal-like cells. **A.** Multilineage organoids demonstrated a reduction in spontaneous beating throughout culture. **B.** Representative calcium flux traces at day 30 and 80 of culture. **C.** Cardiomyocytes in multilineage organoids display mature calcium flux features by showing higher amplitude, Vmax upstroke and downstroke (Conventional vs. Multilineage, **p<0.01, **p<0.005, ***p<0.001, ****p<0.0001). **D.** Cardiomyocytes’ ability to respond to electrical stimuli throughout culture time. Black dashed line: electric stimuli applied**. E.** Representative immunofluorescence images of dissociated cardiomyocytes from conventional microtissues and multilineage organoids. **F.** Cardiomyocyte structural feature metrics. **G.** Cardiomyocyte sarcomere classification (from class 1—immature, undefined sarcomeres—to class 4—mature, parallel, striped sarcomeres). **H.** Day-100 multilineage organoid cardiomyocyte sarcomeres (i), intercalated disks (ii, arrow heads) and glycogen reservoir ultrastructure (iii). **I.** Electrophysiological characteristics of cardiomyocyte subtypes. **J.** Percentage of ventricular and atrial/nodal cardiomyocytes dissociated from conventional microtissues and multilineage organoids. **K.** Representative electrophysiological traces of ventricular and atrial/nodal cardiomyocytes in multilineage organoids at day 30 of culture.

Morphological analysis of dissociated tissues revealed greater structural maturation of individual cardiomyocytes from day-30 multilineage organoids than conventional microtissues, as determined by greater surface area and perimeter, elongated morphology, and highly-aligned and defined sarcomeres **(Figure 4E-G)** (Lundy *et al.*, 2013). More elongated cardiomyocytes with highly-aligned and defined sarcomeres were isolated from day-80 multilineage organoids, reflecting additional structural maturation over time **(Figure 4F,G)**. Electron microscopic imaging of cardiomyocytes from day-100 multilineage organoids revealed a series of ultrastructural features typical of the third-trimester human fetal heart, including: 1) myofibrils closely packed with defined striations and no apparent M bands **(Figure 4H,i)**; 2) cylindrical-shaped mitochondria aligned between the myofibrils **(Figure 4H,ii)**; and 3) large pools of glycogen vesicles, the major energy source during fetal heart development **(Figure 4H,iii)** (Kim *et al.*, 1992;Racca *et al.*, 2016). This advanced degree of iPSC-derived cardiomyocyte structural features is not commonly observed without external stimuli (mechanical, electrical or chemical), thus highlighting the intrinsic ability of the multilineage organoid model to promote cellular maturation of cardiac muscle cells.

We next evaluated cardiomyocyte function using patch-clamp analysis of dissociated single cells, considering multiple action potential parameters (APD90/APD50 ratio, APD30-40/APD70-80 ratio, MDP, peak, amplitude, frequency, and dV/dTMax). Electrophysiology analysis revealed a higher proportion of atrial and nodal-like (atrial/nodal) cardiomyocytes (day 30: 73%) than found in conventional microtissues (day 30: 57%) **(Figure 4 I,J)**. In addition, the number of atrial/nodal cardiomyocytes increased over time (multilineage organoids day 30: 73%, day 60: 80%) **(Figure 4 I,J)**, consistent with the assignment of 98% of cardiomyocytes as atrial-like cells by the *in vivo* cell classifier at day 100 **(Figure 3H; S3G)**. These results reflect the ability of multilineage organoids to specify and mature atrial/nodal cardiomyocytes, which has been challenging to achieve by alternative 3D differentiation methods.

The primary differences between the multilineage and conventional media are the concentrations of ascorbic acid (AA) and calcium. Previous studies have demonstrated that these two components can influence cardiomyocyte differentiation and maturation, respectively(Cao *et al.*, 2012;Wheelwright *et al.*, 2018). To assess the potential impact of different culture medium components alone on cardiomyocyte specification and maturation, patch-clamp analyses were performed in iPSC-cardiomyocytes differentiated in monolayers in conventional cardiac media (Ca^2+^= 0.42mM), multilineage media (AA=100μg/ml, Ca^2+^=1.05mM) and conventional cardiac media supplemented with AA and calcium (AA=100μg/ml, Ca^2+^=0.63mM). The AA- and Ca^2+^-supplemented conventional cardiac media enhanced ventricular cardiomyocyte maturation by promoting an increase in peak, amplitude and upstroke velocity (V/s) of the action potentials, but did not promote the emergence of atrial/nodal cardiomyocyte subtypes (no atrial/nodal cells were identified) **(Figure S5C-E)**. Cardiomyocytes differentiated in multilineage medium (in 2D monolayers) exhibited comparable maturation to those in the supplemented conventional media, but yielded mainly ventricular-like cells (97%) **(Figure S5C-E)**. Therefore, although the media components promoted functional cardiomyocyte maturation, they alone were not sufficient to direct atrial/nodal specification. Furthermore, these experiments suggest that the co-development of the gut tissue, which is absent in conventional cardiac microtissues and 2D monolayers, appears to be a critical factor responsible for enrichment of atrial/nodal cardiomyocytes (>90% at day 100; **Figure 3H, S3J**) and the physiological maturation of cardiac tissue in multilineage organoids to an equivalent level of third-trimester human fetal hearts.

### Multilineage organoids express cardiac and gut master regulatory genes

Many iPSC differentiation protocols suffer from inherent variability in consistency (Volpato and Webber, 2020;Matthys *et al.*, 2020). In our system, when multilineage organoids failed to develop, spheroids instead yielded either solely cardiac microtissues or gut organoids **(Figure 5A,B; S6)**. Despite batch-to-batch differences in spheroid phenotype, all spheroids (n>200) within a single batch derived from the same initial differentiation of progenitor cells manifested identical morphologies and cellularity **(Figure 5A)**. Thus, we hypothesized that subtle differences in the cellular composition of the initial 2D populations of differentiating cells were responsible for the phenotypic outcomes of the spheroids.

**Figure 5.**
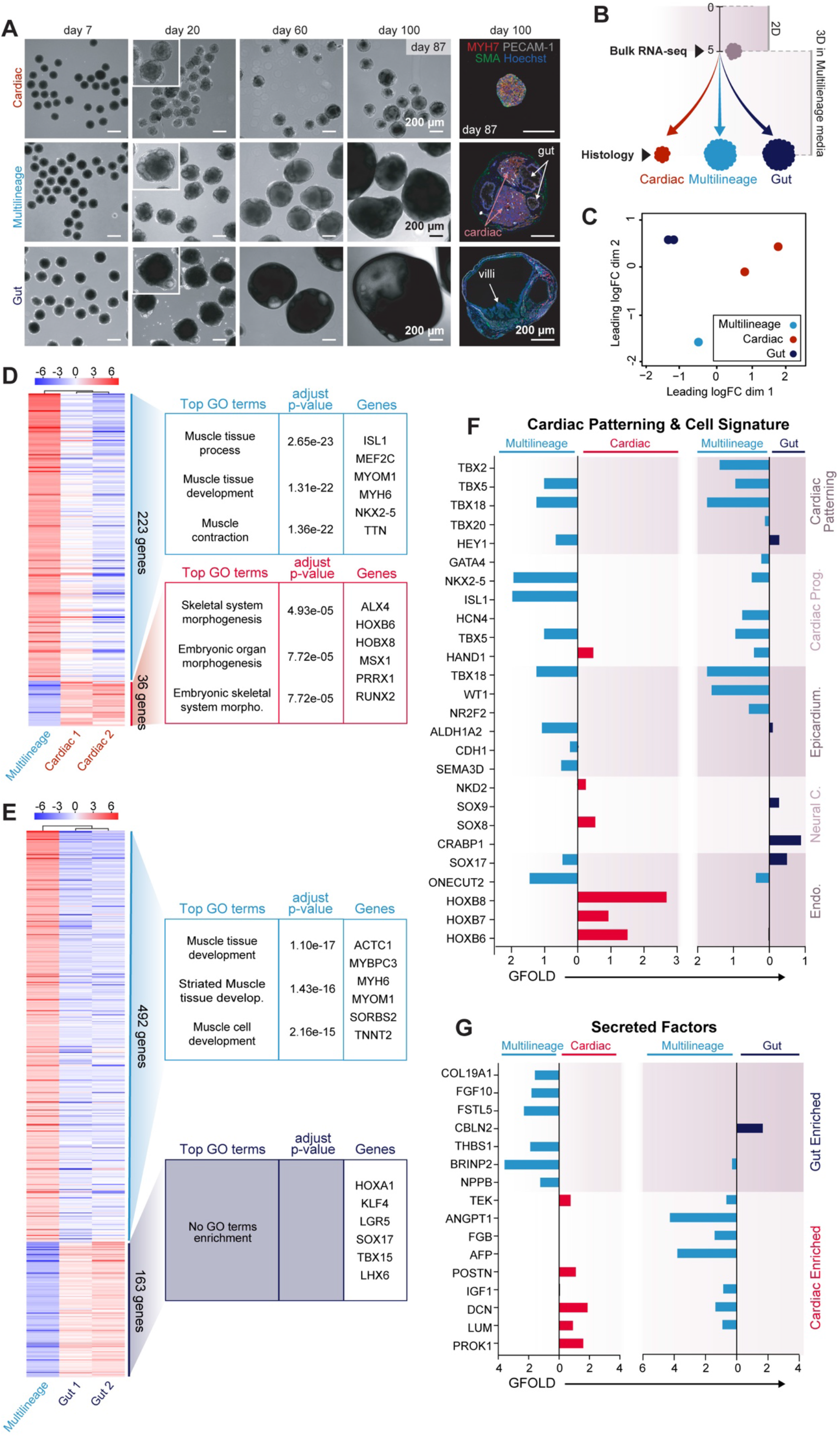
Multilineage organoid progenitor cells have a transcriptomic profile that supports the co-development of cardiac and gut tissues. **A.** Mesendoderm progenitors cultured in AA medium sometimes failed to develop into multilineage organoids, and instead yielded cardiac microtissues or gut organoids. Brightfield and histological analysis revealed phenotypic diversity between these outcomes. **B.** Experimental work flow schematic. **C.** Multidimensional scaling (MDS) plot describes similarity between the progenitor cells that give rise to cardiac microtissues and multilineage and gut organoids. **D.** Heat map of the pairwise comparison of multilineage organoids and cardiac microtissues progenitors evidence a gene transcription profile that favors muscle tissue formation. **E.** Heat map of the pairwise comparison of multilineage organoid and gut organoid progenitors and respective GO terms and genes demonstrated that multilineage progenitor gene expression supports cardiac tissue formation. **F-G.** Targeted pairwise comparison for cell type-specific transcription factor (TF) genes (F) and secreted factor genes (G) in multilineage vs. cardiac (left) and multilineage vs. gut (right) progenitor cells. The multilineage organoid progenitors expressed key heart and endoderm-related TFs and secreted factors highly expressed in gut organoid and cardiac microtissues progenitors.

To evaluate the variability between the starting progenitor populations that gave rise to cardiac microtissues, multilineage organoids, or gut organoids, we performed retrospective analysis of the initial progenitor cell population at day 5 of differentiation by bulk RNA-sequencing, immediately prior to spheroid formation **(Figure 5B)**. Multidimensional scaling (MDS) analysis of the progenitor transcriptomes revealed that different batches had distinct gene expression profiles, and that the progenitors of multilineage organoids represented a unique transcriptional state, roughly equidistant from both cardiac and gut progenitors **(Figure 5C)**. Comparative analysis between cardiac microtissue and gut organoid progenitors indicated that cardiac microtissue progenitors displayed a muscle development-restricted gene expression program. In contrast, progenitors of gut organoids revealed activation of a broader embryonic developmental program, including forebrain, heart, visual and sensory system development **(Figure S7B)**. To dissect the specific transcriptomic features enabling multilineage organoid formation, differential expression analysis was performed to compare the multilineage transcriptional profile to gut organoid and cardiac microtissue progenitors independently. Compared to cardiac microtissue progenitors, multilineage organoid progenitors were enriched in genes related to muscle tissue development and function (*MYH6*, *MYOM1*, *TTN*), as well as transcription factors (TFs) expressed by cardiomyocytes (*MEF2C*, *NKX2-5*, *ISL1*) **(Figure 5D)**, suggesting an increased cardiac signature in the multilineage progenitors. Compared to gut organoid progenitors, multilineage progenitors exhibited a gene expression enrichment of the muscle tissue development program (*ACTC1*, *MYH6*, *MYOM1*) **(Figure 5E)**. No gene ontology term enrichment was observed for the gut organoid progenitors relative to the multilineage progenitors, presumably due to the fact that: 1) the population of gut progenitors express markers of early cardiac differentiation similar to the multilineage progenitors, 2) at early time points (before 20 days), gut organoids phenotypically resemble multilineage spheroids, and 3) and gut tissue development required at least 2 months of 3D culture to manifest clearly with pronounced villi only evident after 100 days **(Figure 5A, S6, S7A; Movie S8)**. Histological analysis revealed that primitive gut structures emerged earlier in the gut organoids **(Figure S7A, day 12)**, however similar structures were not seen in multilineage organoids until later time points **(Figure 5A, day 20-30; Movie S8)**. Thus, the accelerated development of primitive cystic structures in gut organoids might have disrupted the cardiac morphogenesis originally present in multilineage organoids.

In order to dissect specific gene expression differences between multilineage and cardiac or gut progenitors, a targeted pairwise analysis was performed for heart patterning transcription factors, cardiac progenitors and endoderm-related genes. Multilineage progenitors expressed higher levels of T-box (TBX) transcription factors (*TBX2, TBX5, TBX18, TBX20*) that regulate the developing heart(Stennard and Harvey, 2005), in comparison with the gut progenitors, revealing a broader activation of the cardiac patterning program. Notably, in contrast to the cardiac progenitors, multilineage progenitors expressed higher levels of TBX genes related to sinoatrial node formation (*TBX18*) and chamber formation and septation (*TBX5*), in addition to *HEY1,* a gene whose expression is restricted to the atria in the context of the heart(Rutenberg *et al.*, 2006), suggesting a transcriptomic profile more consistent with atrial/nodal cardiomyocyte specification. Furthermore, multilineage progenitors expressed several TFs expressed by cardiac cells (*ISL1*, *NKX2-5*, *TBX5*) and epicardial-related genes (*TBX18*, *CDH1, ALDH1A2, SEMA3D*) at higher levels than cardiac progenitors **(Figure 5F, top)**, consistent with the greater cardiac cell diversity observed in the multilineage organoids as compared to the cardiac microtissues **(Figure 5A)**. Progenitors yielding multilineage organoids also expressed genes involved in gut tube development (*SOX17*, *ONECUT2*), although at lesser levels than gut organoid progenitors (notably for *SOX17*), suggesting that the multilineage populations contained a smaller number or proportion of gut progenitors **(Figure 5F, bottom)**. In total, multilineage organoid progenitors displayed a mixed transcriptomic profile of cardiac and gut progenitors, which could account for their ability to give rise to organoids consisting of both heart and gut cells.

### Multilineage organoid progenitors express cardiac and gut relevant paracrine factor genes

Distinct endoderm subtypes are known to play a role in cardiomyocyte formation through paracrine signaling mechanisms (Holtzinger *et al.*, 2010;Brown and Badylak, 2014;Anderson *et al.*, 2016). To examine potential intercellular paracrine signaling affecting the co-emergent populations, we compared the expression of genes associated with secreted factors between the different progenitor populations **(Figure 5G)**. We first generated a list of secreted factors that were differentially enriched in the cardiac or gut progenitors **(Figure S7D)**. FGF10 and NPPB were identified among the gut-enriched factors. FGF10 is a growth factor that enhances cardiogenic specification, fetal cardiomyocyte proliferation, and has been identified as a second heart field (SHF)-specific marker, indicating a role in heart tissue development (Al Alam *et al.*, 2015;Rochais *et al.*, 2014;Kelly *et al.*, 2001;Chan *et al.*, 2010). NPPB is highly expressed during cardiogenesis, particularly in the differentiating chamber myocardium (Sergeeva and Christoffels, 2013). However, FGF10 and NPPB also promote intestinal cell proliferation, induction of goblet cell differentiation and homeostasis of mesenteric and intestinal tissue(Al Alam *et al.*, 2015;Rochais *et al.*, 2014;Kelly *et al.*, 2001;Sergeeva and Christoffels, 2013). Thus, FGF10 and NPPB could enhance aspects of both cardiac and gut tissue formation. Indeed, multilineage progenitors expressed higher levels of FGF10 and NPPB than the cardiac tissue progenitors **(Figure 5G, top; S7E, left)**. Multilineage progenitors also expressed higher levels of cardiac-enriched factors than the gut progenitors, such as the growth factors IGF1 and ANGPT1, and the ECM proteins DCN and LUM **(Figure 5G, bottom; S7E, right)**. IGF1 enhances mesodermal cell differentiation and subsequent cardiac progenitor cell formation (Engels *et al.*, 2013), and ANGPT1 supports proper atrium chamber morphogenesis (Kim *et al.*, 2018). DCN and LUM, matrix molecule genes enriched in the multilineage progenitors, contribute to the structure and mechanics of the heart by binding to core matrix proteins like collagen I and III, and facilitate thickening of collagen fibrils, respectively, thereby impacting collagen fibrillogenesis (Chakravarti, 2002;Ferdous *et al.*, 2007). Hence, multilineage progenitors expressed increased levels of secreted factor genes relevant to both cardiac and gut tissue morphogenesis compared to either individual tissue alone.

In summary, this retrospective analysis revealed that, in addition to the emergence of cardiac cells within the multilineage organoids, the co-development of gut alongside cardiac cells potentially contributes to more structurally and functionally mature cardiac tissue via the secretion of key cardio-inductive molecules.

## Discussion

Although organoids can recapitulate many physiological properties of tissue development *in vitro*, they typically lack the cooperative paracrine interactions between adjacent tissues that characterize embryogenesis, as well as the diverse multilineage progenitors essential for the development of complex organs such as the heart. Current strategies to increase organoid complexity have focused on the fusion of differentiated organoids (Koike *et al.*, 2019;Bagley *et al.*, 2017), but these approaches miss the interactions between germ layers that occur during the early steps of embryogenesis to initiate the formation of different organs. Although a couple of recent reports demonstrate the ability to generate primitive cardiac tissues coincident with endoderm derivatives from pluripotent stem cells with greater fidelity than traditional embryoid body differentiation (Rossi *et al.*, 2020;Drakhlis *et al.*, 2021), such organoids consist of many cell types from different germ lineages. However, in contrast, here we describe a multilineage organoid model system in which only cardiac (mesoderm) and gut (endoderm) tissues emerge in a sequential manner. This multilineage organoid recapitulates morphogenic events that lead to the emergence of key structural features of the developing human heart and gut. Past studies of the interactions between heart and gut during development indicated that endoderm-derived cells generate paracrine signals and mechanical forces that support heart formation (Rudy-Reil and Lough, 2004;Brown *et al.*, 2010;Uosaki *et al.*, 2012;Sugi and Lough, 1995;Kidokoro *et al.*, 2018),(Hosseini *et al.*, 2017;Kelly *et al.*, 2001;Drakhlis *et al.*, 2021;Rossi *et al.*, 2020). Consistent with those findings, our multilineage organoid also demonstrates that endoderm-derived cells can positively impact cardiomyocyte phenotype, suggesting a broad contribution of endoderm-derived cells to heart morphogenesis.

We generated multilineage organoids by aggregating early committed mesendoderm progenitors derived from human iPSCs in permissive media conditions. The sequential emergence of functional cardiac and gut-like tissues mimicked human embryonic morphogenesis, in which the first heart beat (3 weeks of gestation) is followed one week later by the extension of the gut tube into fore-, mid- and hind-gut structures (Hill, 2020b;Hill, 2020a). At early stages of *in vitro* culture, multilineage organoids resemble human fetal heart formation (~5 weeks of gestation), evidenced by the formation of a surface layer of epicardial cells and a core of cardiomyocytes. Similar features were also observed in conventional microtissues, derived from progenitors spheroids cultured in cardiac-permissive medium. However, only multilineage organoids exhibited significant growth and preserved the exterior epicardial layer over the long term, as well as, a preferential differentiation into atrial/nodal cells, in stark contrast to other cardiac organoid models that also contain endoderm-derived cells(Rossi *et al.*, 2020;Drakhlis *et al.*, 2021). The epicardium is an important source of IGF2, a potent cardiac mitogen involved in cardiac wall thickening (Shen *et al.*, 2015;Perez-Pomares and De La Pompa, 2011). It also generates cardiac fibroblasts and smooth muscle cells (SMCs) that invade the heart following an epithelial-to-mesenchymal transition (Smits *et al.*, 2018). Similarly, in our organoid model, we observed cardiomyocyte expansion, as well as the later emergence of SMCs and endothelial cells (which occurs at >5 weeks of gestation in human fetal heart)(Smits *et al.*, 2018;Hirakow, 1983). We propose that the co-emergence of multiple key progenitor cells contributed to the cellular and structural complexity of the cardiac compartment of the multilineage organoids.

The presence of gut tissue developing alongside the cardiac tissue improved the phenotypic and functional maturation of the organoid cardiomyocytes, whose structural features resembled those of native third-trimester human fetal heart. To date, more mature features of cardiomyocytes have only been achieved *in vitro* by subjecting early contractile iPSC-derived cardiomyocytes on a 3D collagen gel to an increasing regimen of electrical stimulation (Ronaldson-Bouchard *et al.*, 2018). Additionally, the prevailing view of atrial and ventricular cardiomyocyte specification is that the different cardiomyocyte subtypes derive from distinct mesoderm populations (Lee *et al.*, 2017;Devine *et al.*, 2014). Our organoid model shows that the emergence of cardiomyocytes with an atrial/nodal phenotype occurs in parallel with the formation of primitive gut: whereas at day 20, multilineage organoids contained equivalent amounts of ventricular (MLC2V+) and atrial/nodal (MLC2V-) cardiomyocytes, by day 30, when primitive mid-hindgut (MHG) structures emerged, the proportion of atrial/nodal cardiomyocytes was greater **(Figure 2 D,E)**. Therefore, we speculate that similar to other endoderm subtypes MHG endoderm secretes factors that impact the development of the heart, and specifically promotes atrial/nodal specification. Indeed, previous studies have shown that other endoderm derivatives, such as anterior visceral endoderm and anterior foregut, promote cardiac mesoderm specification and formation of new cardiomyocytes through the secretion of cardiac-inductive factors, like Activin A and FGF2 (Sugi and Lough, 1995;Mummery *et al.*, 2003;Anderson *et al.*, 2016;Lough and Sugi, 2000). FGF2 is also expressed by the epithelial, stromal, and SMCs in the intestine, but its effects on gut development have not been fully defined (Dignass and Sturm, 2001;El-Hariry *et al.*, 1997). Our multilineage organoid model of cardiac and gut co-emergence offers a novel platform to investigate how specific neighboring tissues synergistically interact to promote co-development and functional maturation.

The complexity of the intestinal tissue in multilineage organoids mimics that of the small intestinal mucosa that forms adjacent to enteric SMCs. The formation of an SMC layer in close proximity to the gut epithelium in our multilineage organoids is notable since SMCs have so far only been observed in gut organoids after *in vivo* transplantation (Munera *et al.*, 2017). Furthermore, the SMCs in multilineage organoids exhibited spontaneous peristaltic-like contractility, which has only been observed in fully mature cells *in vivo*, or in *in vitro* cultures of primary cells together with stromal cells and gut pacemaker cells (interstitial cells of Cajal) seeded onto a scaffold (Gays *et al.*, 2017;Kobayashi *et al.*, 2018). Other gut organoid models form an intestine-like mucosa only upon embedding in an exogenous extracellular matrix (i.e. Matrigel) (Munera *et al.*, 2017;Sato *et al.*, 2009). Thus, our organoid model creates a microenvironment permissive for gut tissue formation in the absence of exogenous extracellular matrix components or morphogenic signaling cues, likely due, in part, to the enrichment of ECM components (e.g. glycosaminoglycans and hyaluronic acid) detected in the sub-epicardial/ peritoneum-like region where the gut epithelium clusters arise.

While multilineage organoids arose robustly from all the aggregated spheroids within a single experiment, not all differentiation experiments gave rise to multilineage organoids. In those alternative instances, we obtained gut organoids or cardiac microtissues instead, at similar frequencies, from the same initial aggregation and culture conditions. Gene expression analysis of the starting populations that yielded the different phenotypic outcomes indicated that multilineage organoids result from progenitors with enriched expression of heart-related patterning TFs, cardiomyocyte contractile machinery related genes, and SOX17, a definitive endoderm master regulator gene. By contrast, cardiac microtissue progenitors expressed a transcriptome that was less enriched for heart cell related genes, and gut organoid progenitors strongly expressed the endoderm marker, SOX17. The expression of endoderm-specific genes at the progenitor stage suggests that not all cells undergo the intended cardiac mesoderm commitment, and that the subsequent emergence of intestinal tissue in the multilineage and gut organoids results from the permissive culture medium supporting the expansion and differentiation of the endoderm progenitors. The analysis of genes encoding secreted molecules in the gut organoid progenitors revealed high expression of FGF10, which functions as an autocrine and paracrine signaling molecule in heart development as well as in the gut, where it regulates the balance between goblet and Paneth cells (Al Alam *et al.*, 2015;Rochais *et al.*, 2014;Kelly *et al.*, 2001). Our multilineage organoid model can be useful for the future dissection of paracrine interactions between cardiac and gut tissues, and identification of specific factors that promote morphogenesis and maturation of adjacent tissues.

Future efforts can also focus on more robustly controlling the proportion of mesoderm and endoderm cells in the initial progenitor pool. By modulating the expression of cardiac and gut regulatory genes in distinct sub-populations of progenitors, it should be possible to reduce batch-to-batch variability when generating multilineage organoids. Genetic engineering approaches have been demonstrated to control cellular patterning and subsequent adoption of divergent cell fates – fundamental first steps to regulate phenotypic co-emergence and morphogenesis of human iPSC-derived organoids(Libby *et al.*, 2019;Libby *et al.*, 2018;Mackay *et al.*, 2014). In addition, we envisage that “mosaic” transcriptional modulation approaches can be expanded for the development of multilineage organoids that support the co-emergence of distinct tissues other than heart and gut.

Overall, our model represents a new class of organoids characterized by the co-development and maturation of two distinct tissues arising from a mixed pool of mesendoderm progenitors. This organoid model opens new avenues to study genetic diseases that affect both the heart and the gut that have been challenging to investigate, such as chronic atrial and intestinal dysrhythmias (Chetaille *et al.*, 2014). Organoids have been primarily developed with the intent to model individual tissues in isolation (Munera *et al.*, 2017;Takasato *et al.*, 2016), and to date only limited attempts have been made to generate more complex and heterotypic organoids, mostly via fusion of pre-differentiated cells and/or organoid tissues (Xiang *et al.*, 2017;Koike *et al.*, 2019;Bagley *et al.*, 2017). In contrast, multilineage organoids demonstrate that two specific tissues from distinct germ lineages can arise in parallel within a single spheroid to synergistically support each other’s development, and thus will be useful to advance the understanding of cell or tissue interactions during early tissue specification, structural and functional maturation, and/or disease development. Although they may not mimic all aspects of human development, multilineage organoids do offer a more direct view into tissue cross-talk mechanisms than alternative *in vitro* systems or animal models currently in use. Multilineage organoids represent a logical next step in the generation of more physiological *in vitro* models of human development, and therefore broaden the possibilities for modeling early embryogenesis or complex multi-organ disorders *ex vivo*.

## Acknowledgments

The authors would like to thank the Stem Cell Core (Dr. Po-Lin So), Histology and Light Microscopy Core (Dr. Meredith Calvert) and Genomics Core (Dr. Natasha Carli and Jim McGuire) at Gladstone Institutes, and Electron Microscopy Lab (Dr. Danielle Jorgens) at UC Berkeley for all the help and expertise shared. The authors thank Dr. Bruce Conklin’s laboratory for providing WTC11 GCaMP iPS cell line, Dr. Deepak Srivastava’s laboratory for providing antibodies against MYL2 and MYL3, and Dr. Katja Schenke-Layland for providing human fetal heart paraffin blocks. A special thanks to Drs. Tracy Hookway, Irfan Kathiriya, Sanjeev Ranade and Elphège Nora for critical discussions of this work. The authors also acknowledge the Gladstone Scientific Editing Department (Drs. Francoise Chanut and Kathryn Claiborn) for assistance with manuscript editing. This work was supported by the California Institute of Regenerative Medicine Grant (LA1_C14-08015) and the Gladstone BioFulcrum Heart Failure Research Program.

## Author contributions

A.C.S. and T.C.M. conceived the study, interpreted the data, and wrote the manuscript. A.C.S. performed iPSC differentiations, microtissues formation and maintenance, histology studies, preparation of the cells for patch clamp analysis, calcium imaging, bulk and single-cell RNA-seq experiments and scRNA-seq analysis. O.B.M. acquired and analyzed calcium flux analysis, assisted on microtissues maintenance. M.H.L. performed patch clamp data acquisition and analysis. D.A.J. developed the python script for calcium flux and patch clamp analysis, performed the alignment of single-cell and bulk RNA-seq data and assisted with the single-cell and bulk RNA-seq analysis. M.A.K. and V.N. prepared bulk RNA-seq libraries and assisted in microtissue maintenance. D.T. performed light sheet microscopy and analysis. A.P.B. performed the *in vivo* cell classifier analysis and with B.G.B. contributed to interpretation of the results. M.A. and B.G.B. assisted with bulk RNA-seq data analysis and B.G.B. contributed to experimental design and interpretation of results. O.B.M., D.A.J., M.A.K, V.N., M.A., and B.G.B. contributed to manuscript editing and discussion.

## Materials and Methods

### Human induced pluripotent stem cell culture

WTC11 human induced pluripotent stem cells (iPSCs) genetically modified with fast green fluorescent calcium indicator GCaMP6f (Huebsch *et al.*, 2015;Mandegar *et al.*, 2016) were maintained on tissue culture plates coated with 80μg/mL growth factor-reduced (GFR) Matrigel (Corning) and fed daily with mTeSR1 medium (StemCell Technologies). When the iPSCs reached ~70% confluency, about every 3 days, they were dissociated to single cells using Accutase (Innovative Cell Technologies) and re-plated at density of 1×10^4^ cells/cm^2^ in mTeSR1 medium containing 10μM Y-27632 ROCK inhibitor (ROCKi; Selleckchem) for 24h.

### Differentiation to mesendoderm-derived progenitors

GCaMP-WTC11 iPSCs were differentiated to mesendoderm-derived progenitor cells following a chemically-defined, serum-free protocol modulating Wnt/β-catenin signaling (Lian *et al.*, 2012). Briefly, iPSCs were seeded at a density of 3×10^4^ cells/cm^2^ and fed with mTeSR1 medium until 100% confluency was reached (~3 days; termed differentiation day 0). At day 0, cells were fed with RPMI1640 medium with B27 Supplement minus insulin (denoted as RPMI/B27-; Life Technologies) supplemented with 12μM CHIR-99021 (Selleckchem). Exactly 24h later, the medium was changed to fresh RPMI/B27-. On day 3, cells were fed with RPMI/B27-supplemented with 5μM IWP2 (Sellekchem) for 48h. On day 5, cells were either dissociated for tissue formation or fed with RPMI/B27-medium.

### Mesendoderm progenitor cell spheroid formation and culture

Mesendoderm progenitor cells were dissociated at differentiation day 5 with Accutase and aggregated as previous described (Hookway *et al.*, 2016). Briefly, dissociated cells in RPMI/B27-medium with 10μM ROCKi were seeded into 800μm pyramidal agarose wells at a density of 2.4×10^4^ cells/well. After 24h, cell spheroids were removed from the agarose wells by gently pipetting using wide bore tips. Spheroids were maintained in rotary suspension culture at a density of 65 spheroids/mL in 6-well low-attachment tissue culture plates (Hookway *et al.*, 2016). On day 7 of the differentiation protocol, 2 days after tissue aggregation, medium was changed to either conventional cardiac-permissive medium, RPMI/B27+ (RPMI1640 medium with B27 supplement with insulin; Life Technologies)(Lian *et al.*, 2012) or epicardial-permissive medium (Advanced DMEM/F12 (Gibco) with 100μM L-Ascorbic Acid (Sigma))(Bao *et al.*, 2016) and refreshed every 3 days thereafter.

### Calcium imaging and functional analysis

Calcium flux was analyzed by recording fluorescence of GCaMP signal in the spheroids. At day 30, 40, 50, 60, 70, 80, 90*, 100*, and 365*, conventional and multilineage microtissues (n = 4-17 per condition) were equilibrated in Tyrode’s solution (137 mM NaCl, 2.7 mM KCl, 1 mM MgCl_2_, 0.2 mM Na_2_HPO_4_, 12 mM NaHCO_3_, 5.5 mM d-glucose, and 1.8 mM CaCl_2_; Sigma-Aldrich) for at least 30min at 37°C and then imaged on a Zeiss Axio Observer Z1 inverted microscope equipped with a Hamamatsu ORCA-Flash 4.0 sCMOS camera. Electrical field stimulation between 0.5Hz and 8Hz was applied with the MyoPacer (IonOptix) to evaluate functional properties and eliminate differences in intrinsic beat rate between individual spheroids. Calcium transients were recorded using Zen Professional software (v.2.0.0.0) with 10ms exposure and 100 frames per second. Circular regions of interest (ROI) with a diameter of 59.52 pixels were selected in the beating core of the spheroids and the mean fluorescence intensity of the calcium transients was plotted against time. Additional metrics of calcium handling properties, such as amplitude, stroke velocities, and beat rate, were analyzed using a custom Python-script. Source code is available at https://github.com/david-a-joy/multilineage-organoid (made public upon acceptance). *Of note, conventional samples were not evaluated at these timepoints, because they were only viable until day 80 of culture.

### Calcium imaging with light sheet microscopy

In order to assess calcium activity at higher resolution, light sheet fluorescence microscopy (LSFM) was used to optically section live multilineage organoids and conventional microtissues and record calcium transients *in situ* (Turaga *et al.*, 2020). Organoids/microtissues were suspended in 1.5% low-melt agarose made up in Tyrode’s solution in size 2 glass capillaries (~1mm inner diameter; Zeiss). The Zeiss z.1 light sheet microscope used for imaging was equipped with two PCO.edge sCMOS cameras, 10x 0.2 NA illumination lens, 20x 1.0 NA detection lens, and 488nm laser, and the live samples were maintained at 37°C. 3D calcium imaging stacks were acquired at ~120ms per frame and each z-spacing was 0.86μm per frame. Single optical sections (3-4μm light thickness) were excited at 488nm and imaged at ~20Hz for 52 seconds (1000 total frames).

### Patch clamp analysis

Conventional microtissues (n = 10 per condition), and multilineage organoids (n = 5-10 per condition), as well as conventional 2D differentiated cardiomyocytes, were washed in PBS and dissociated by successive 20min incubations in 0.25% Trypsin at 37°C (1 incubation for monolayers and maximum of 5 incubations for tissues) until complete dissociation to single cells. Dissociated cells at a density of 0.03×10^6^cells/cm^2^ were seeded onto GFR Matrigel-coated glass coverslips for patch-clamp recordings in medium containing 10μM ROCKi. 24h after seeding, cells were kept in culture media without ROCKi until the experiments began. To conduct experiments, coverslips containing hiPSC-CMs were transferred to a heated RC-26 recording chamber (Warner Instruments), mounted to a Nikon inverted microscope and continuously perfused with warmed Tyrode’s solution. Experiments were performed at 35 ± 1°C. Action potentials were recorded from spontaneously beating cardiomyocytes using the perforated patch technique with borosillicate glass pipettes (Sutter Instruments) with typical resistance of 2-4 MΩ. The Tyrode’s solution consisted of (in mM): 140 NaCl, 5.4 KCl, 1.8 CaCl_2_, 1 MgCl_2_, 10 glucose, and 10 HEPES, with the pH adjusted to 7.4 with NaOH. The patch pipette solution consisted of (in mM): 150 KCl, 5 NaCl, 5 MgATP, 10 HEPES, 5 EGTA, 2 CaCl_2_, and 240 μg/mL amphotericin B, with the pH adjusted to 7.2 with KOH. Data were acquired in current clamp mode using an Axopatch 200B amplifier and Digidata 1440A digitizer coupled to pClamp 10 software (Molecular Devices). Data were digitized at 10 kHz and filtered at 1 kHz. Action potential parameters were determined using Clampfit 10 software within pClamp. hiPSC-derived cardiomyocytes were classified as either ventricular or nodal/atrial subtype after considering multiple action potential parameters, including the APD90/APD50 ratio, APD30-40/APD70-80 ratio, MDP, peak, amplitude, frequency, and dV/dTMax (Ma *et al.*, 2011;Churko *et al.*, 2018).

### Cardiomyocyte structure analysis (area, perimeter, AR, circularity and sarcomere score)

Day-30 and 80 conventional* (n =10 per condition) and multilineage microtissues (n = 5-10 per condition) were dissociated into single cells as described above (patch clamp analysis sections) and seeded onto GFR Matrigel-coated glass coverslips. 24h post-seeding, the cells were fed with fresh medium without ROCKi for at least 48h to allow the cells to acquire their natural morphology. Coverslips were then fixed in 4% paraformaldehyde (EMS), stained with the cardiomyocyte-specific marker cardiac Troponin T (cTnT), labeled with Hoechst to detect nuclei, and mounted on glass slides using ProLongΩ Gold Antifade Mountant (ThermoFisher Scientific). Representative images were acquired for each sample condition using an inverted microscope (Zeiss Axio Observer Z1). For each cell, area, perimeter, circularity, and aspect ratio (AR) were quantified using Fiji software. In addition, each cardiomyocyte was assigned a score for sarcomere development between 1 and 4, defined as: 1) unaligned sarcomeres and absence of a striped pattern; 2) continuous sarcomeres, well ordered but not in parallel; 3) continuous sarcomeres, organized mostly in parallel, but with less-defined striped pattern; 4) continuous sarcomeres with striped pattern, organized mostly in parallel **(Figure M1)** (Nguyen *et al.*, 2014;Judge *et al.*, 2017). The percentage of cells in each class of sarcomere development was calculated per sample and timepoint. * Of note, conventional samples were not evaluated at day 80, because only a few microtissues were available and viable at this later stage.

**Figure M1.**
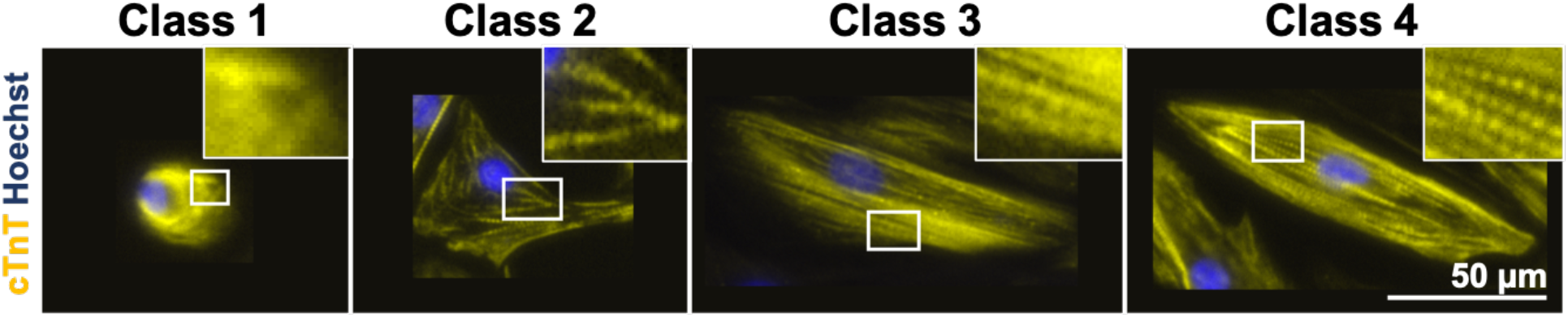
Representative images of the different classes of sarcomere development.

### Histology, immunostaining, and in situ hybridization

Conventional microtissues (n=10 per condition), cardiac organoids (n=10 per condition), multilineage organoids (n=5-10 per condition), and gut (n=5-10 per condition) organoids were fixed in 10% neutral buffered formalin (VWR) with 0.03% eosin (Newcomer Supply) at room temperature for 1h and washed with PBS. Samples were embedded in HistoGel Specimen Processing Gel (ThermoFisher Scientific), paraffin processed and embedded, cut into 5μm sections, and adhered to positively charged glass slides. Slides were then deparaffinized with xylene, re-hydrated through a series of decreasing ethanol concentrations (100%, 100%, 95%, 80%, 70%), and rinsed in tap water. Hematoxylin and eosin (H&E), Russell-Movat Pentachrome, and Safranin O stainings were performed according to manufacturer’s instructions and the slides were mounted with Cytoseal 60 (Richard-Allan Scientific) and glass coverslips.

For immunofluorescence staining, heat-induced epitope retrieval was performed by immersing slides in Tris-EDTA buffer (Cellgro) pH 9.0 or citrate buffer (Vector Laboratories) pH 6.0 in a 95°C water bath for 35min. Slides were cooled for 20min at RT and washed 3x with PBS. Samples were permeabilized by slide immersion in 0.2% or 0.5% Triton X-100 (Sigma) in PBS, washed 3x in PBS, and blocked in 1.5% normal donkey serum (NDS; Jackson ImmunoResearch). Primary and secondary antibody cocktails were diluted in blocking solution **(Table S3)**. 3 PBS washes were performed after primary (overnight, 4°C) and secondary antibody (1h, RT) incubations. For in situ hybridization analysis, RNAscope Multiplex Fluorescent Reagent Kit v2 was used on paraffin sections, following user manual #323100-USM. Human RNAscope probes used in this study were: WT1-C2 (catalog #415581); FOXA2-C1 (catalog #832221); TBX18-C3 (catalog #832231). Nuclei were counter-labeled with Hoechst and coverslips were mounted on slides using ProLong™ Gold Antifade Mountant. Samples were imaged on the Zeiss Axio Observer Z1.

### Whole-mount immunostaining

Conventional microtissues (n=5) and multilineage organoids (n=5) were fixed in 4% paraformaldehyde at room temperature for 1h with agitation (40 rpm). Samples were then washed 3x in stain buffer (2% NDS, 0.1% Tween-20 (Sigma-Aldrich) in PBS) and permeabilized for 30min in 1.5% Triton X-100, followed by a 3h incubation in blocking buffer (3.5% NDS, 0.1% Tween-20 in PBS). Microtissues/organoids were incubated with TBX18 and cTnT primary antibodies overnight at 4°C and secondary antibodies plus phalloidin and Hoechst stains for 4h at RT **(Table S3)**, followed by 3 final washes in stain buffer prior to imaging.

### Whole-spheroid structural analysis by light sheet microscopy

Whole-mount immunostained conventional microtissues and multilineage organoids were suspended in 1.5% low-melt agarose (made up in PBS; IBI Scientific) in size 2 glass capillaries immediately prior to imaging. Microtissues/organoids were each imaged at three angles (120° rotations between views), and then stitched with multi-view reconstruction to provide isotropic resolution throughout the sample, as previously described (Turaga *et al.*, 2020). Volumetric reconstruction and size analyses were performed using Imaris 8.3 software.

### Single-cell RNA sequencing

Multilineage and gut organoids at day 100 of culture (n = 5-8) were washed in PBS and dissociated by successive 20-min incubations in 0.25% trypsin at 37°C (maximum of 5 incubations) until complete dissociation to single cells. 10,000 cells in PBS supplemented with 0.08% FBS were prepared for sequencing by droplet encapsulation using the Chromium Controller and library preparation with the 10x Genomics Single Cell 3’ v2 Library and Gel Bead Kit (10x Genomics). Libraries were sequenced using the NovaSeq S4 (Illumina) to a minimum depth of 50,000 mean reads per cell. Sequences were demultiplexed and aligned to human reference genome grch38 using *CellRanger* v3.0.2, and downstream analysis was performed with Seurat v.3.1. Cells with fewer than 1,500 genes were excluded from analysis and cell identity was assigned to each cluster by cross-referencing cluster marker genes with known cardiac and gut markers. Cell-cycle did not introduce heterogeneity in scRNA-seq data. Raw data is available at Geo under the accession number GSE153075 (access token: qvslsuecdpofjup).

### Classification of In Vivo Cardiac Cell Types

To evaluate the transcriptomic similarity between multilineage organoid heart cells and human fetal heart cells, an *in vivo* heart cell classifier was applied. A multiclass logistic regression (lr) model was trained on *in vivo* cardiac cell types(Asp *et al.*, 2019). To have consistent features, we ran SCTransform (default, version 3.1.5.9) independently on each dataset and then took an intersection of their top 3,000 highly variable genes. A total of 1,418 genes, transformed as Pearson residuals, were used to train the model. The resulting expression matrix of 3,717 single cells from a 6.5–7 post conception week (PCW) heart with established cardiac cell types was used to train the model using a ‘sag’ solver and cross-entropy loss cost function with the Python package sklearn (version 0.22.2.post1).

A two-dimensional UMAP (version 0.4.4) latent space was fit to *in vivo* subepicardial cells, fibroblast-like, smooth muscle cells/fibroblast-like, and atrial cardiomyocytes. We used the lr model’s feature space for UMAP dimensionality reduction. A sklearn C-support vector classifier was then trained on the four *in vivo* cell types using the UMAP latent space. *In vitro* cells that the lr model classified as one of the four *in vivo* cell types, with probabilities greater than or equal to eighty-five percent, were transformed into the UMAP latent space. The trained C-support vector classifier was then applied to the *in vitro* cell’s UMAP transform embedding. The Seaborn (version 0.10.1) library was used to visualize UMAP transformations.

### Bulk RNA sequencing

To reduce batch effects between RNA-sequencing analysis of the mesendoderm-derived progenitors between experiments, bulk RNA-sequencing was applied. Day 5 progenitor cells that gave rise to cardiac (n=2), multilineage (n=1), or gut organoids (n=2) were lysed with Trizol (Ambion) and RNA was extracted using the Direct-zol Miniprep kit (ZymoResearch) and quantified using the NanoDrop 2000c (ThermoFisher Scientific). RNA-seq libraries were created using the SMARTer Stranded Total RNA Sample Prep Kit (Takara Bio) and sequenced using NextSeq500/550 High Output v2.5 kit (spiked with 10% PhiX; Illumina) to a minimum depth of 29 million reads per sample. The sequences were aligned to grch38 using HiSat2 (Kim *et al.*, 2019), reads were quantified using featureCounts (Liao *et al.*, 2014), and differential expression was determined using GFOLD(Feng *et al.*, 2012). Relative gene expression level was determined using the “l” normalization algorithm (Zhang *et al.*, 2019). Raw data is available at Geo under the accession number GSE153071 (access token: utwxqsawhjcpfgz).

### Statistical analysis

All results were obtained in three independent experiments, unless otherwise stated, and shown as mean ± standard error of the mean (SEM). The statistical analysis of the data was performed using Prism 8 with statistical significance determined at p < 0.05. Tukey’s multiple comparison test was applied. Whenever normality was verified through D'Agostino-Pearson omnibus normality test, the t-Student test was applied and the Independent Samples Mann-Whitney U test was used for the remaining analyses. A blinded approach (labeling samples with an alphanumeric code) was implemented for analyzing cardiomyocyte morphological features (area, perimeter, AR, circularity, sarcomere scoring), patch clamp, histology, and immunostaining.

---

**Figure S1.**
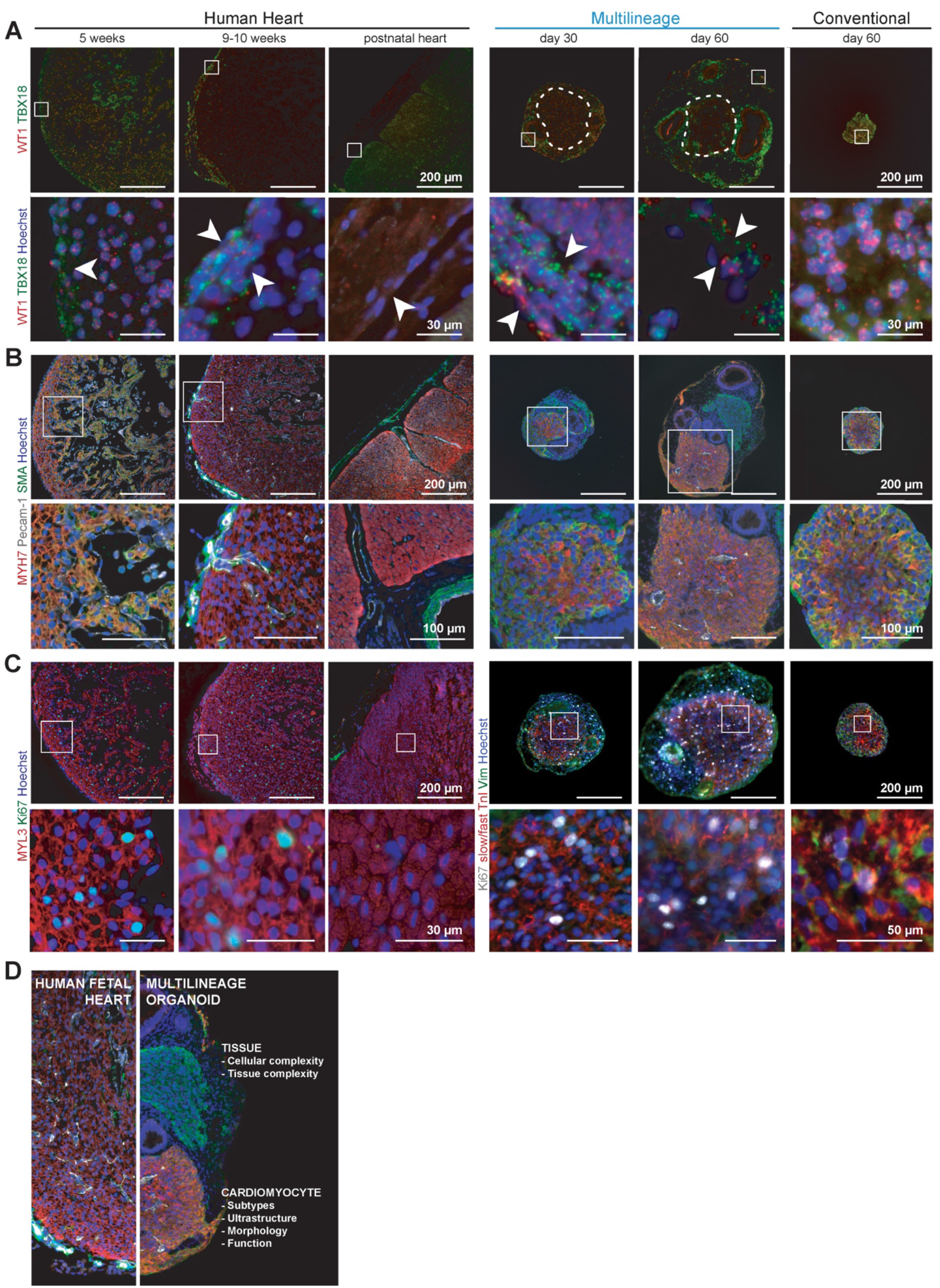
Related to Figure 1 and 2. Comparison of conventional and multilineage microtissues with human heart development (**A-B**). **A.** RNAscope validation of TBX18 and WT1 transcripts in native human heart tissue, multilineage organoids, and conventional microtissues. Dashed line denotes boundary of the cardiac core. **B.** Identification of cardiomyocytes (MYH7+SMA-, MYH7+SMA+), smooth muscle cells (SMA+MYH7-), and endothelial cells (Pecam-1+) in native human heart tissue, multilineage organoids, and conventional microtissues. **C.** Proliferating cardiomyocytes in human ventricle wall tissue (MYL3+Ki67+), multilineage organoids, and conventional microtissues (slow/fast TnI+Ki67+). **D.** Comparative images exhibiting similarities between the cardiac compartment of native human fetal heart tissue and that of the multilineage organoids.

**Figure S2.**
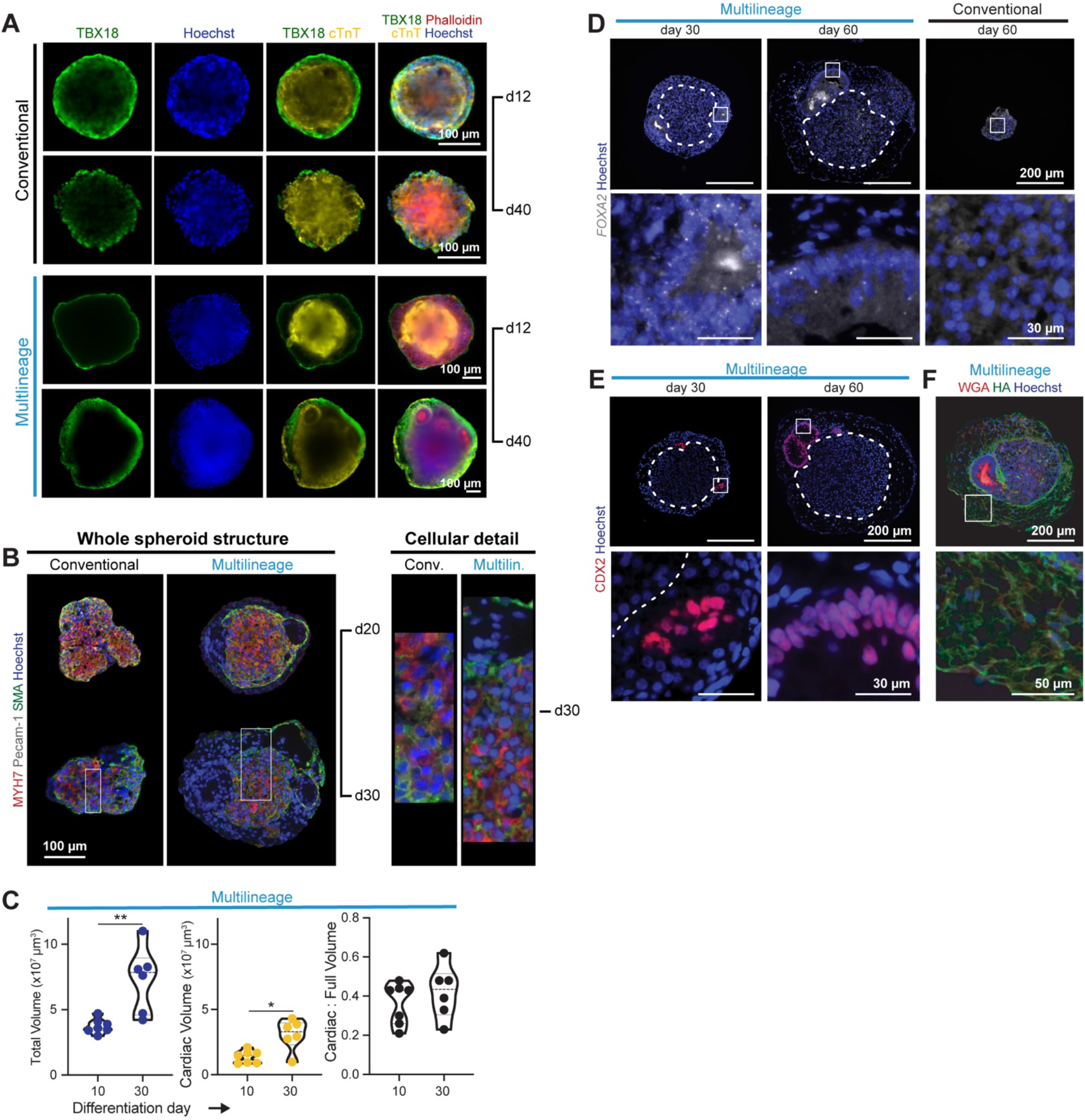
Related to Figure 1 and 2. Multilineage organoid histological and volumetric analysis. **A.** Detailed light-sheet images of the epicardial layer (TBX18+) of conventional and multilineage microtissues. **B.** Conventional and multilineage microtissues demonstrate the presence of cardiomyocytes (MYH7+), smooth muscle cells (SMA+) and MYH7+/SMA+ cardiomyocytes (d20, more immature). **C.** Total volume (blue), cardiac core volume (yellow), and ratio of cardiac:total volume (black) of the multilineage organoids at days 10 and 30 of culture. *p<0.05, **p<0.005. **D-E.** Epithelial-like structures in multilineage organoids correspond to primitive midgut based on the detection of FOXA2 transcript by RNAscope (D) (absent in conventional microtissues) and CDX2 protein by immunostaining (E). **F.** Hyaluronic acid (HA) is expressed in the interstitial tissue (tissue surrounding the cardiac core and the gut) in the multilineage organoids. White squares regions in high magnification at the bottom panels. Dashed line boundary of the cardiac core and gut.

**Figure S3.**
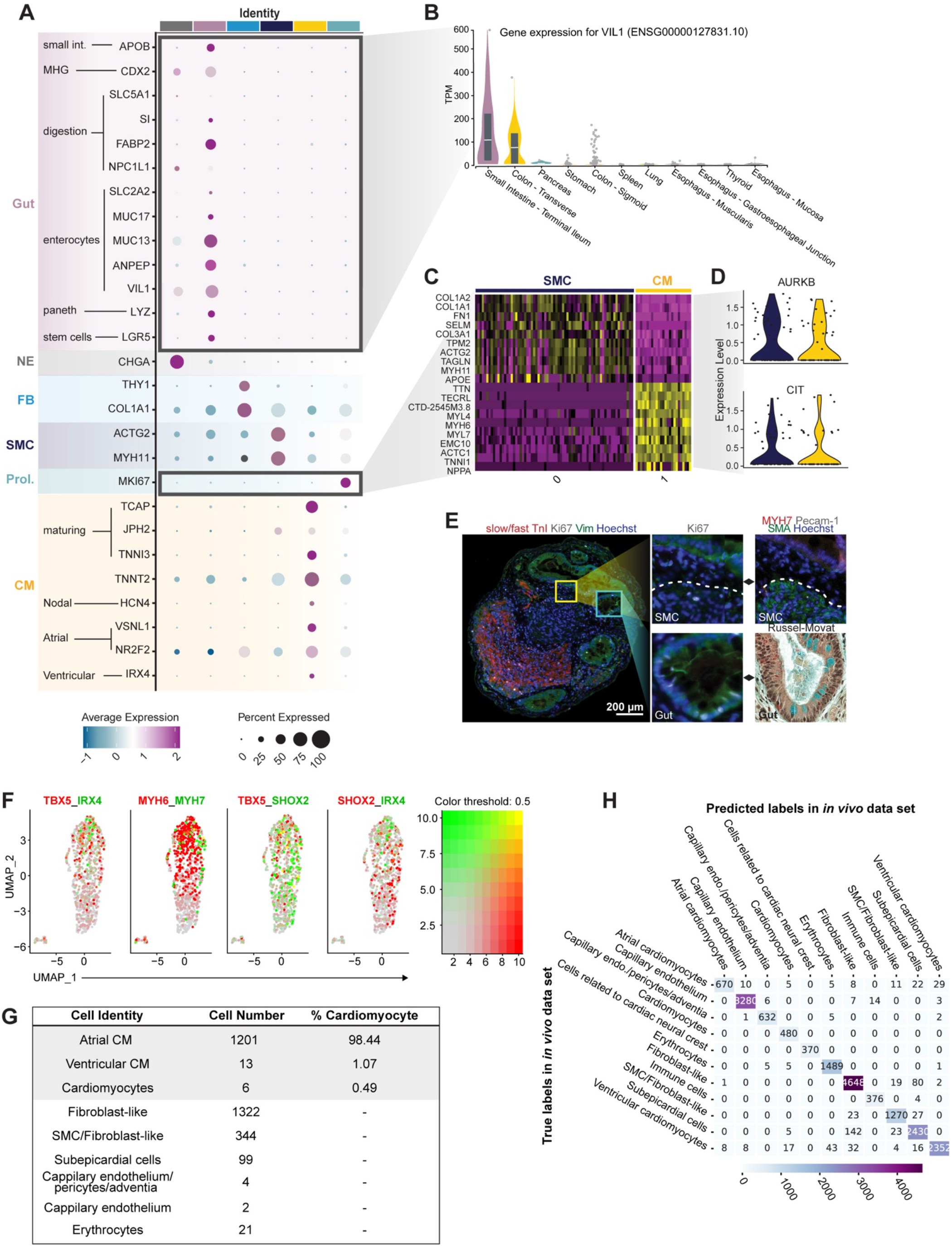
Related to Figure 3. Transcriptional characterization of day-100 multilineage organoids. **A.** Dotplot of key genes that define cell type identity of UMAP clusters as cardiomyocytes (CM), proliferating cells (Prol.), smooth muscle cells (SMC), fibroblasts (FB), enteroendocrine cells (EnteroE), and gut. **B.** Violin plot of VIL1 expression across different endoderm-derived tissues using GTEx tissue dataset (https://www.gtexportal.org/). **C.** Heatmap of secondary analysis of re-clustered proliferating cells identified SMCs (cluster 0) and CMs (cluster 1) as the main proliferative cell types at day 100. **D.** Violin plot of key karyokinesis-related genes in proliferating SMCs and CMs. **E.** Histological validation of SMC proliferation (yellow square; ki67+, SMA+) and gut cell proliferation (blue square; ki67+, Russel-Movat) in a day-100 multilineage organoid. Dashed line denotes SMC region. **F.** Feature plot showing co-expression of key atrial, ventricular and pacemaker related genes in day-100 organoids. **G.** Table showing the distribution of organoid heart cells classified to various cell identities by the *in vivo* cell classifier. **H.** Confusion matrix comparing test vs. predicted cell type labels for human developing fetal cardiac cells(Asp *et al.*, 2019).

**Figure S4.**
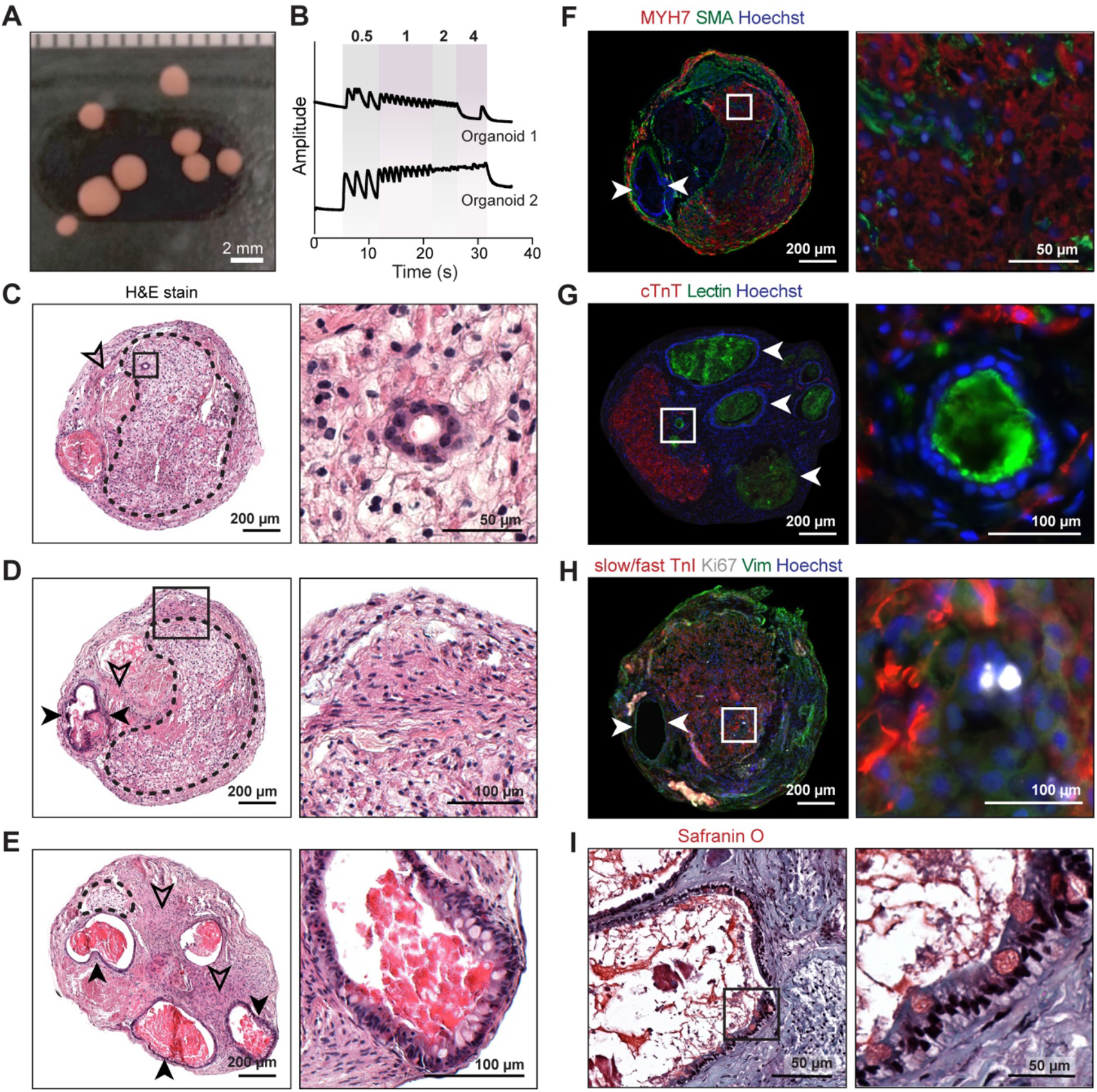
Related to figure 2. 1 year-old multilineage organoids. **A.** Macroscopic field of view of multiple 1 year-old organoids. **B.** Calcium handling response to electrical stimulation (0.5-4Hz). **C-E.** Hematoxylin and eosin stain (H&E) reveals vasculature-like structures (**C**), smooth muscle cells (**D**), and intestine (**E**). Dashed line denotes cardiac core; solid arrowhead indicates gut; outlined arrowhead indicates smooth muscle cells. **F.** Immunostaining highlights the presence of cardiomyocytes (MYH7+) and smooth muscle cells (SMA+). **G.** Lectin-positive vasculature-like structures are found close to the cardiac compartment (inset) and non-specific lectin expression was also identified in gut structures (solid arrowheads). **H.** Identification of actively proliferating cells by Ki67+ immunostaining. **I.** Safranin O staining reveals presence of goblet cells (counterstained in red) in the gut structures.

**Figure S5.**
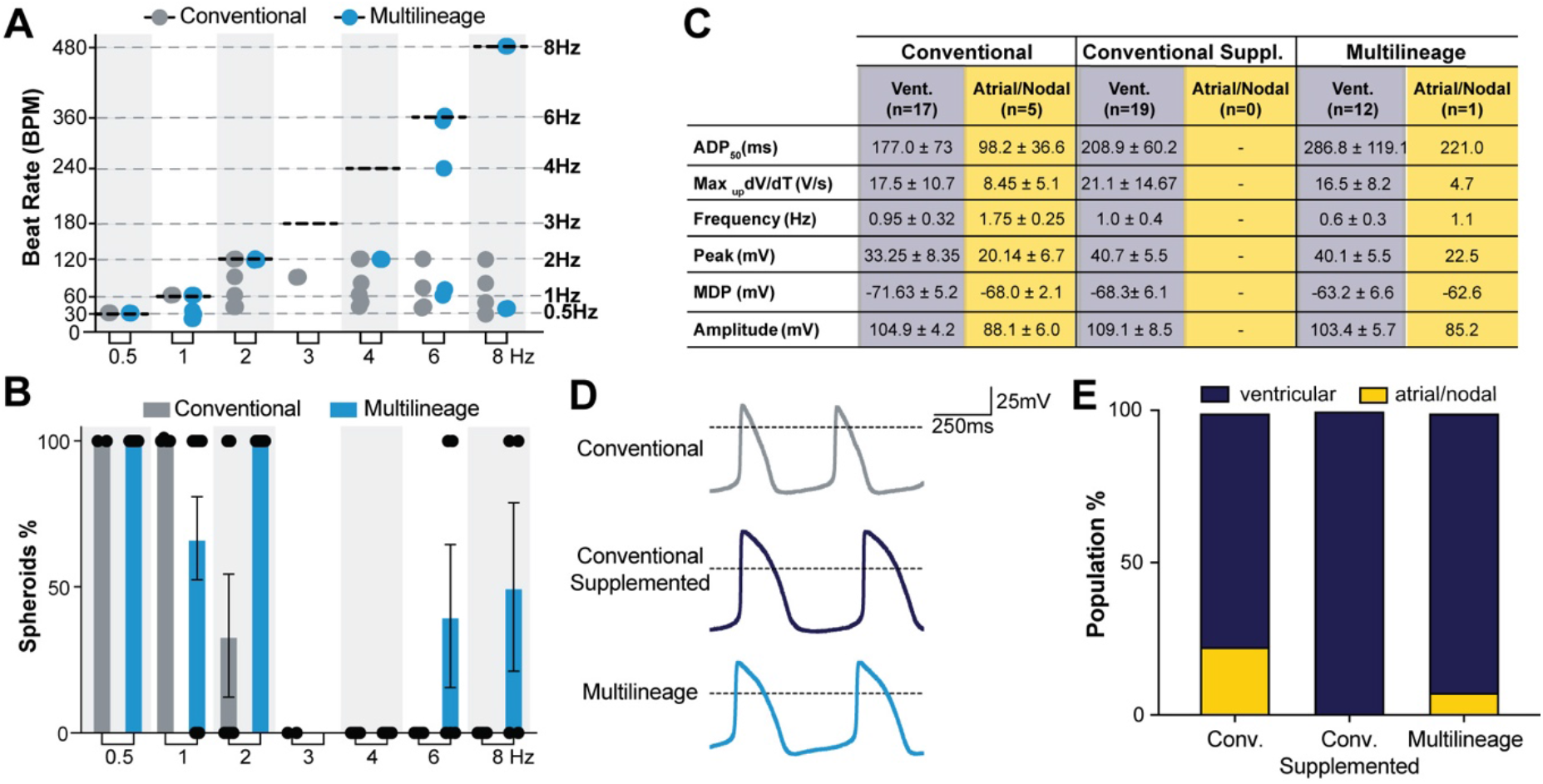
Related to Figure 4. Cardiomyocyte functional analyses. **A.** Calcium handling response of day 80 conventional and multilineage spheroids subjected to electrical field stimulation at increasing frequencies (0.5-8Hz). **B**. Percentage of conventional and multilineage spheroids capable of responding to the electrical field (0.5-8Hz) stimulation applied at day 80 of culture. **C.** Table displaying electrophysiological properties of day-30 cardiomyocytes differentiated in 2D in various media conditions: conventional media (RPMI/B27+, Ca^2+^=0.42mM), conventional media supplemented with ascorbic acid (AA) and calcium (Conventional Suppl., AA=100 μg/ml, Ca^2+^=0.63mM) and multilineage-permissive media (AA=100 μg/ml, Ca^2+^=1.05mM). **D.** Representative traces of ventricular cardiomyocytes in the different media conditions. **E.** Percentage of ventricular and atrial/nodal cardiomyocytes across the different media conditions.

**Figure S6.**
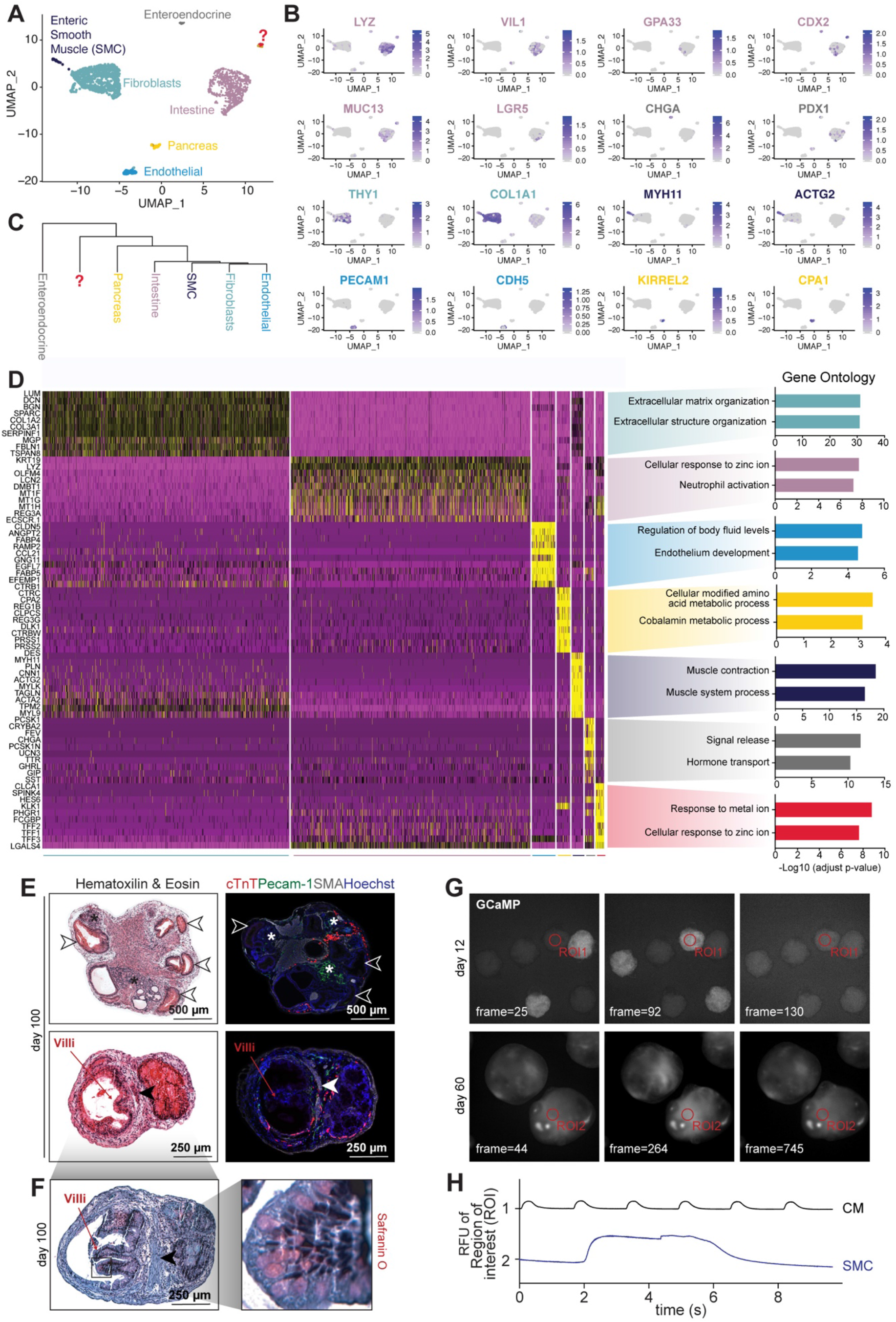
Related to Figure 5. Analysis of day-100 gut organoids by single-cell RNA sequencing, histology, and immunostaining. **A.** UMAP plot colored by cell identity. **B.** Feature plot of key genes used to assign cell-type identity to UMAP clusters. **C.** Hierarchical clustering of the different UMAP cell populations. **D.** Heatmap of top 10 differentially expressed genes that define each UMAP cluster with associated biological process gene ontology analysis. **E-F.** Histological analysis of gut organoids from 2 independent experiments. Hematoxylin & eosin staining (left, E), immunostaining to identify the presence of endothelial cells (Pecam-1+), smooth muscle cells (SMA+, solid arrowhead), and remnant cardiomyocytes (cTnT+) (right, E) and Safranin O staining to identify the presence of goblet cells (red, F). Outlined arrowhead, intestine tissue; *, pancreas-like tissue. **G-H.** Representative images of spontaneous calcium flux transients (G) in gut organoids at day 12 (top, ROI1) and 60 (bottom, ROI2) of culture and respective ROIs traces (H) (Movie S8).

**Figure S7.**
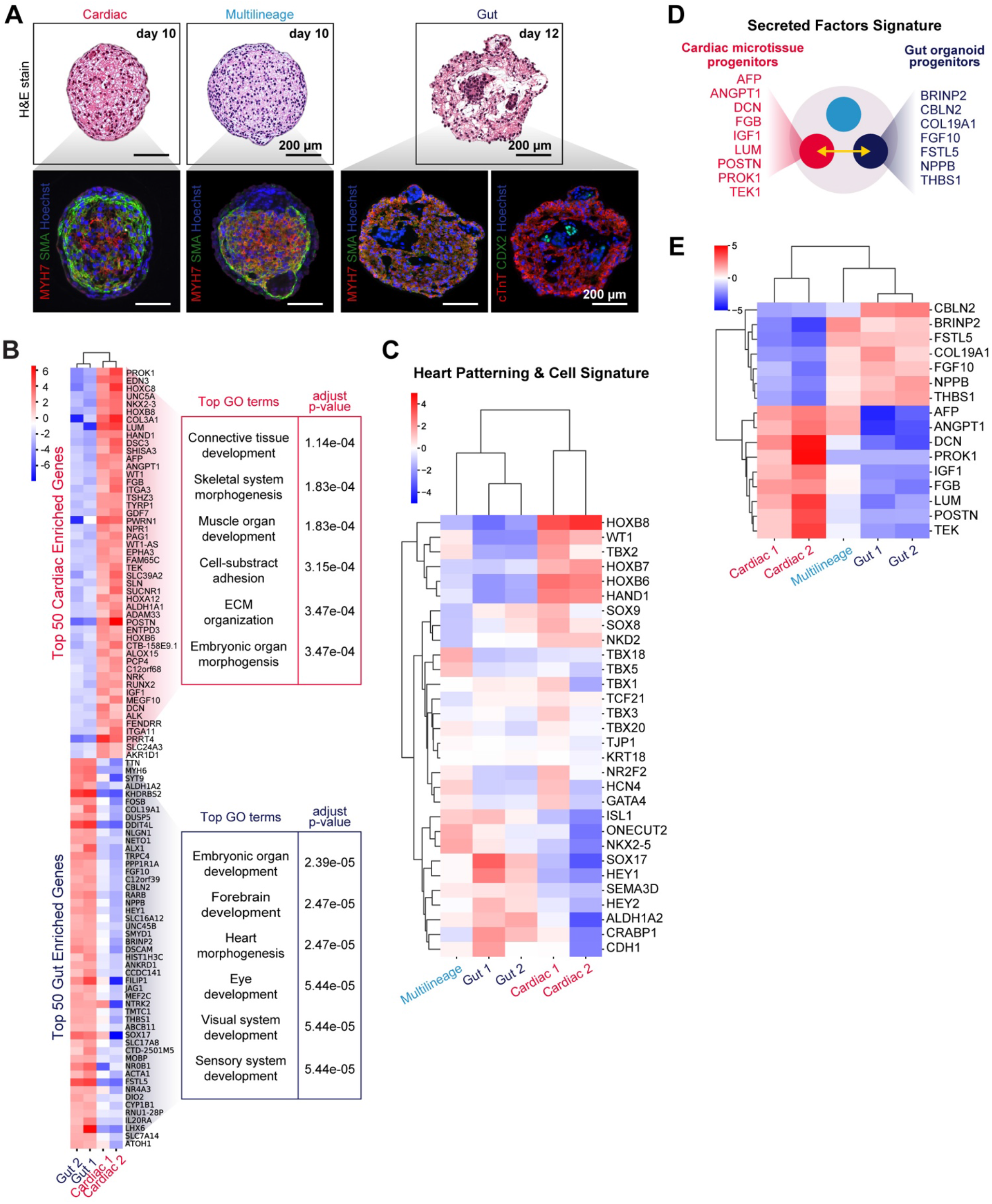
Related to Figure 5. Transcriptional variability of the progenitor populations across experiments. **A.** Spheroid phenotype diversity observed across experiments at early stages of culture. **B.** Heat map and respective gene ontology (GO) terms analysis of cardiac microtissues (top) versus gut organoids progenitors (bottom). **C.** Heatmap of the differentially expressed genes analyzed through a target analysis of relevant cardiac- and gut-related genes in the progenitor cell populations between experiments. **D.** List of highly expressed genes encoding secreted factors in cardiac microtissues and gut organoids. **E.** Heatmap of the differentially expressed genes detailed in (**D**) in the progenitor cell populations for each experiment.

**Table S1.** List of genes that describe each cell cluster identity derived from Seurat analysis.

**Table S2.** *In vivo* cell classifier gene features and weights for each cell type prediction.

**Table S3.** Panel of dyes, primary and secondary antibodies used for whole mount immunostaining, immunocytochemistry and paraffin sections.

| Protein | Source | Catalog # | Dilution |
| --- | --- | --- | --- |
| Alexa Fluor 488 Donkey anti-mouse IgG | Invitrogen |  | 1:400 |
| Alexa Fluor 555 Donkey anti-rabbit IgG | Invitrogen |  | 1:400 |
| Alexa Fluor 647 Donkey anti-goat IgG | Invitrogen |  | 1:400 |
| Alexa Fluor 647 Phalloidin | Invitrogen | A22287 | 1:100 |
| CDX2 | R&D Systems | AF3665 | 1:100 |
| Collagen IV | Millipore | AB769 | 1:100 |
| Ki67 | Invitrogen | Ab15580 | 1:500 |
| Smooth muscle actin | DAKO | M0851 | 1:100 |
| HABP | Sigma-Aldrich | H9910 | 1:65 |
| Hoechst 33342 | Thermo Scientific | 62246 | 1:10000 |
| MYL3 | Abcam | Ab680 | 1:1200 |
| MYH7 | Sigma-Aldrich | HPA001239 | 1:100 |
| MLC2-v | Abcam | Ab79935 | 1:200 |
| PECAM-1 | Santa Cruz | SC-1506 | 1:1000 |
| TBX18 | R&D Systems | MAB63371 | 1:100 |
| To-Pro-3 Iodide | Thermo Fisher | T3605 | 1:5000 |
| Troponin I Slow and Fast | Abcam | ab110132 | 1:100 |
| Troponin T | Abcam | ab45932 | 1:400 |
| Vimentin | Millipore | AB1620 | 1:100 |
| WGA | Thermo Fisher | AF350 | 1:100 |

**Movie S1.** Spontaneous calcium transients, depicted by GCaMP fluorescence signal, of day-30 conventional microtissues.

**Movie S2.** Spontaneous calcium transients of day-30 multilineage microtissues.

**Movie S3.** Representative 3D reconstruction of day-12 multilineage microtissue by light sheet microscopy. Total volume was quantified based on a nuclear label (blue, Hoechst) and volume of the cardiac core was based on cardiomyocyte-specific marker, cardiac troponin T (yellow, cTnT). Stromal-like cell population identified by phalloidin stain (red).

**Movie S4.** Representative 3D reconstruction of day-30 multilineage microtissue by light sheet microscopy, with Hoechst labeling nuclei (blue) and cTnT marking cardiomyocytes (yellow). Stromal-like cell population identified by phalloidin stain (red).

**Movie S5.** Lightsheet microscopy-based acquisition of the rhythmic calcium flux transients of cardiomyocytes in day-30 multilineage organoids in the absence of electrical stimulation.

**Movie S6**. Lightsheet microscopy-based acquisition of the rhythmic and sporadic calcium flux transients from cardiomyocytes and smooth muscle cells, respectively, in day-76 multilineage organoids in the absence of electrical stimulation.

**Movie S7.** Calcium transients of 1-year-old multilineage organoids subjected to electrical stimulation from 0.5-4Hz.

**Movie S8.** Calcium transients of day-12 and day-60 gut organoids.

